# Trained phages constrain evolutionary routes to antibiotic resistance in *Escherichia coli*

**DOI:** 10.64898/2026.08.03.742405

**Authors:** Soffi Kei Kei Law, Christopher Blake, Hock Siew Tan, Michael J. McDonald

**Affiliations:** School of Science, Monash University Malaysia, Jalan Lagoon Selatan, 47500 Subang Jaya, Selangor, Malaysia; School of Biological Sciences, Monash University, Clayton, Victoria, Australia; Monash University Malaysia Genomic Platform, Bandar Sunway, 47500, Selangor Darul Ehsan, Malaysia; Centre to Impact AMR, Monash University, Clayton, Victoria, Australia; ARC Centre for Excellence Centre for Mathematical Analysis of Cellular Systems

**Author notes:** **Corresponding authors:** &.

## Abstract

Bacteriophages are major drivers of bacterial evolution and can impose strong selection on traits that mediate resistance and fitness. This creates the possibility of using phage to steer microbial adaptation toward predictable phenotypic outcomes. Yet bacterial populations often evolve phage resistance through diverse genetic routes, not all of which impose meaningful costs or trade-offs. Such evolutionary flexibility can allow bacteria to escape phage selection without acquiring the desired phenotype, limiting the reliability of phage-based evolutionary steering. Here, we develop a directed phage evolution strategy that eliminates cost-free resistance pathways, forcing bacteria into evolutionary trajectories that restore antibiotic susceptibility. We isolated two novel Microviridae phages (BLS2 and BLS5) that infect *E. coli* BL21 and found that bacteria evolved resistance through two distinct mechanisms: mutations in lipopolysaccharide (LPS) biosynthesis genes, which carried substantial fitness costs and conferred sensitivity to erythromycin, and loss-of-function mutations in *yajC*, which imposed no fitness penalty and maintained full antibiotic resistance. We show that YajC is a previously uncharacterised accessory receptor required for phage genome translocation across the inner membrane. We then used this mechanistic understanding to design a phage training regime that evolved phages capable of infecting *yajC* mutants, thereby closing this cost-free escape route. Bacteria challenged with trained phages were constrained to evolve LPS-based resistance, and were consequently resensitised to erythromycin. Our results demonstrate that directed phage evolution can reshape the fitness landscape of bacterial resistance, channelling adaptation along trajectories that carry predictable phenotypic costs – with broad implications for phage-mediated manipulation of microbial populations.

**Graphical Abstract:** 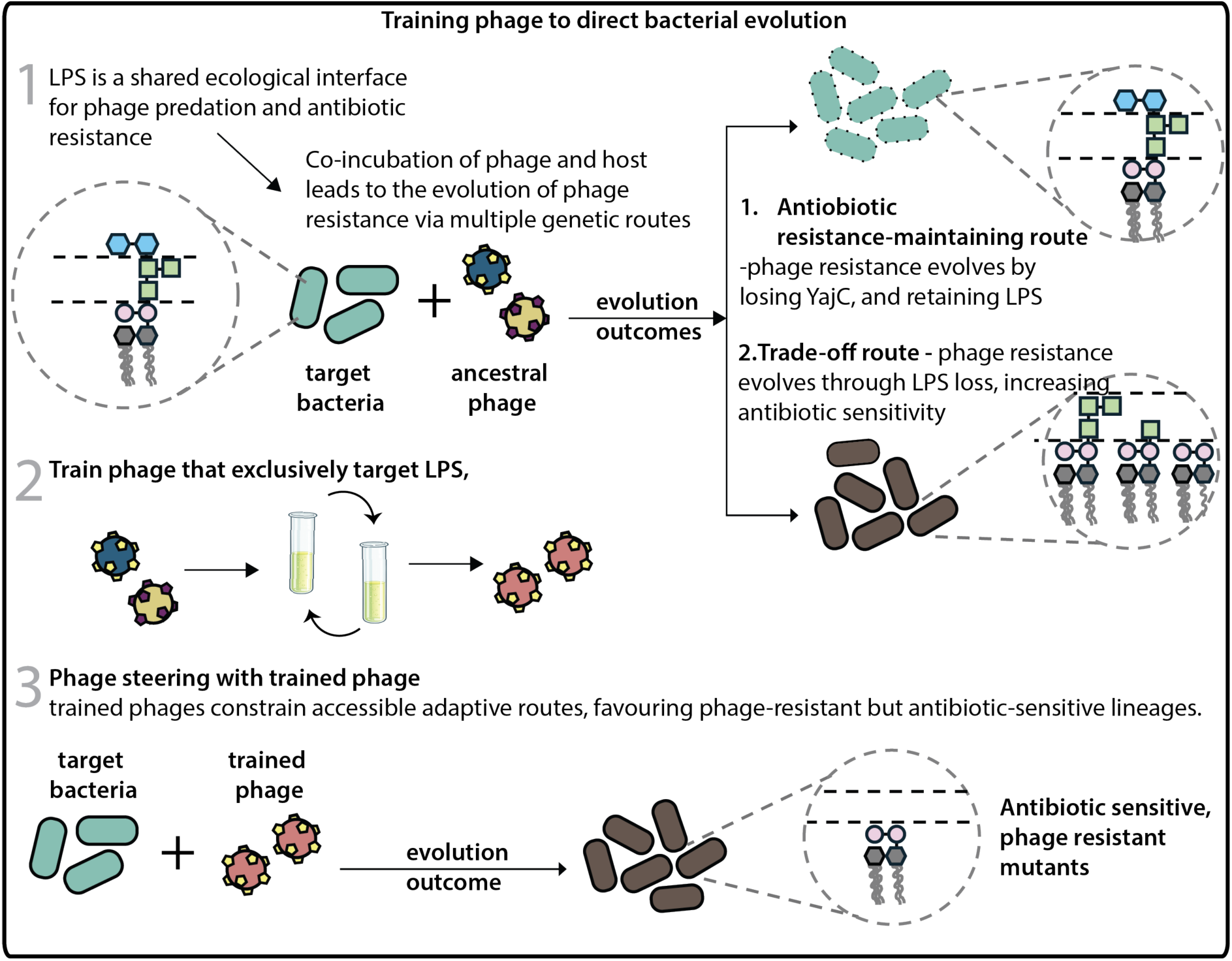

## Introduction

Bacteriophages are the most abundant biological entities on Earth and exert strong selective pressures on bacterial populations. Through cycles of infection, resistance, and counter-adaptation, phage-bacteria coevolution shapes the population dynamics of microbial communities. Understanding and harnessing these coevolutionary dynamics offers opportunities to manipulate microbial populations in ways that go beyond simply killing target bacteria. Phage steering is an approach that exploits trade-offs in bacterial antimicrobial resistance mechanisms to direct bacterial adaptation toward antibiotic sensitivity (*1–4*).

Multiple cases where phage steering could be deployed to reverse antibiotic resistance have been identified. For example, phage targeting *Pseudomonas aeruginosa* drives the evolutionary loss of multidrug efflux pumps, restoring sensitivity to tetracycline (*2*). Similarly, lipopolysaccharide (LPS) targeting phage drives the evolution of antibiotic-sensitive LPS mutants (*3, 5*). However, a fundamental limitation of phage steering as a strategy for directing bacterial evolution is that bacteria rarely have only a single route to phage resistance. In most phage-host systems, resistant mutants arise through diverse genetic changes, and only a subset of these carry the fitness costs or collateral sensitivities that phage steering relies upon(*C-S*). The remaining mutants – those that evolve resistance without paying a phenotypic cost – effectively escape the intended evolutionary trap. This heterogeneity in resistance mechanisms means that even when a beneficial trade-off exists, a significant proportion of the surviving bacterial population may retain resistance to both the phage and the antibiotic, undermining the therapeutic potential of phage steering (*9, 10*).

Phage training involves experimental evolution of bacteriophage populations under controlled conditions to enhance or modify their ability to infect specific bacterial hosts (*11–13*). By propagating phage in the presence of resistant bacterial strains, phage populations can be driven to evolve counter-adaptations — expanding host range, increasing infectivity against resistant variants, or altering receptor usage. Phage training has been deployed to improve phage infectivity against resistant bacterial strains (*14*) and even expand or narrow the host range to target pathogenic bacteria more effectively (*15*). However, the logic of phage training has not yet been extended to the problem of closing cost-free resistance pathways – that is, evolving phage that specifically eliminate the bacterial escape routes that lack fitness trade-offs. In this study, we sought to extend the scope of phage training and to determine whether phages can be trained to block the evolution of mutants that simultaneously confer resistance to both phage and antibiotics.

## Results & Discussion

### *E. coli* BL21 evolves phage resistance by mutating LPS synthesis genes and y*ajC*

To test this idea, we isolated two novel single-stranded DNA phages, *Escherichia* phages BLS2 and BLS5, from wastewater using *E. coli* BL21 as the host (Tables S1-2, Fig. S1A–B). We found that both phages inhibited bacterial growth, but *E. coli* BL21 reliably evolved phage resistance (Fig. S1C – 1D). We sequenced both phage genomes and found only a single synonymous substitution (BLS2 T2536; BLS5 C2536) difference between the two. This mutation rests in an open reading frame (P11, Table S2) in a region highly transcribed in phage phiX174, the best characterised of the Microviridae (*16*).

We isolated and sequenced 19 phage-resistant bacteria (Fig. 1) and found that most (BLS2, 66%; BLS5, 92%) had mutations in genes that encode lipopolysaccharide biosynthesis (Fig. 1, Tables S3 – 4), consistent with previous studies of Microviridae-resistant *E. coli* mutants (*17, 18*). LPS is a critical component of the outer membrane of Gram-negative bacteria (*19*), and the precise impact of LPS gene mutations on LPS composition is well established (*18, 20–23*) (Fig. 1). Based on the mutations identified in our DNA sequencing, we classified the LPS profiles of all the phage-resistant mutants into three commonly used phenotypic categories. In addition, we also identified phage-resistant bacteria with mutations in the non-LPS-related gene *yajC* (Tables S3–4).

**Figure 1:**
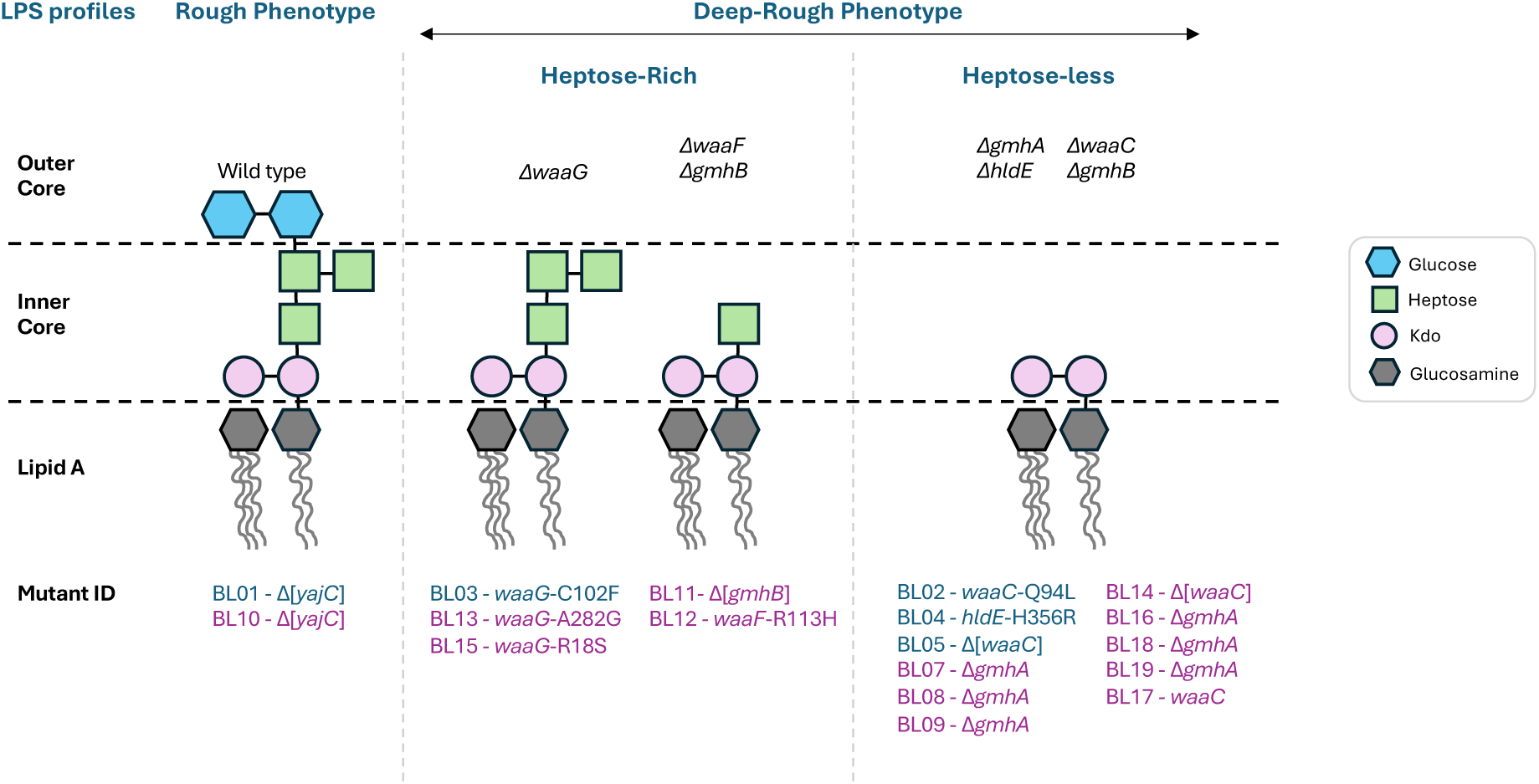
Phage-resistant mutants. Schematic representation of LPS profiles for phage-resistant mutants (blue for phage BLS2 and purple for phage BLS5). The prediction was made based on the complete loss of gene function in the top panel (*Δgene*). Kdo refers to 3-deoxy-D-manno-oct-2-ulosonic acid. “Rough” and “Deep rough” are terms used to refer to the LPS status of a bacterial cell.

### YajC is required for BLS2 and BLS5 infections

YajC is an inner membrane protein that forms part of the Sec protein translocase complex (*24*). Although YajC is upregulated in *E. coli* C when infected with *Microviridae* phage X174 (*25*), its precise function is unknown. Deletion of YajC was reported not to impair the functionality of the bacterial LPS (*26*), despite LPS being critical for maintaining membrane integrity (*27*) and frequently serving as a receptor for phage attachment (*9, 14, 28-31*). These findings suggest that YajC does not directly influence LPS-mediated phage adsorption. To determine if YajC is essential for *Microviridae* infection of *E. coli*, we constructed a *yajC* deletion mutant of *E. coli* BL21 (Fig. S2A). The Δ*yajC* knockout mutant was completely resistant to phage BLS2 (Dunnett’s test, p = 2 x 10^−4^) and phage BLS5 (Dunnett’s test, *p* < 1 x 10^−4^), while complementation with a plasmid borne *yajC* restored sensitivity (BLS2: Dunnett’s test, *p* = 0.56; BLS5: Dunnett’s test, *p* = 0.94) for BLS2 and BLS5, respectively (Fig. S2B – 2C, Tables S6 – 7, Supplementary Data S2C). To test whether YajC was required for intracellular replication and assembly, we transformed the *E. coli* wild type and mutants with naked phage-DNA, circumventing the requirement for phage to adsorb and inject DNA. We found that naked phage DNA was able to establish a productive infection in *E. coli* BL21 Δ*yajC* that was indistinguishable from the wild type and complemented strain, indicating that YajC is dispensable for infection after DNA injection (Fig. 2A, Fig. S3B, Supplementary Note 1). Next, we looked at phage adsorption, finding that phages were able to adsorb to the *yajC* mutant just as well as to the wild type (Fig. 2B, Fig S3A, Supplementary Note 1). Finally, we tested whether YajC contributed by facilitating phage genome delivery. In the tailless microphages, DNA is delivered by the H tube, which penetrates through the outer and inner membrane, channelling the phage genome into the host cytoplasm (*32, 33*). Since YajC is located in the inner membrane, we hypothesised that this protein serves as an accessory receptor that stabilises the H tube during genome translocation, a supposition supported by indirect evidence in the Salmonella targeting Podoviridae phage P22 (*26*). To directly test this idea, we carried out a phage ejection assay, where the phage is incubated and allowed to adsorb to the host (Fig. 2C). If the phage can inject its DNA into the host, the now-empty capsid will become non-viable. If the phage cannot inject its DNA, then phages recovered from the surface of this host will be viable. We found that phage (BLS2) that adsorbed to host cells without YajC were not able to inject their DNA (Fig. 2D), supporting that YajC is required by the wild-type phage for DNA injection (Fig. S3E).

**Figure 2.**
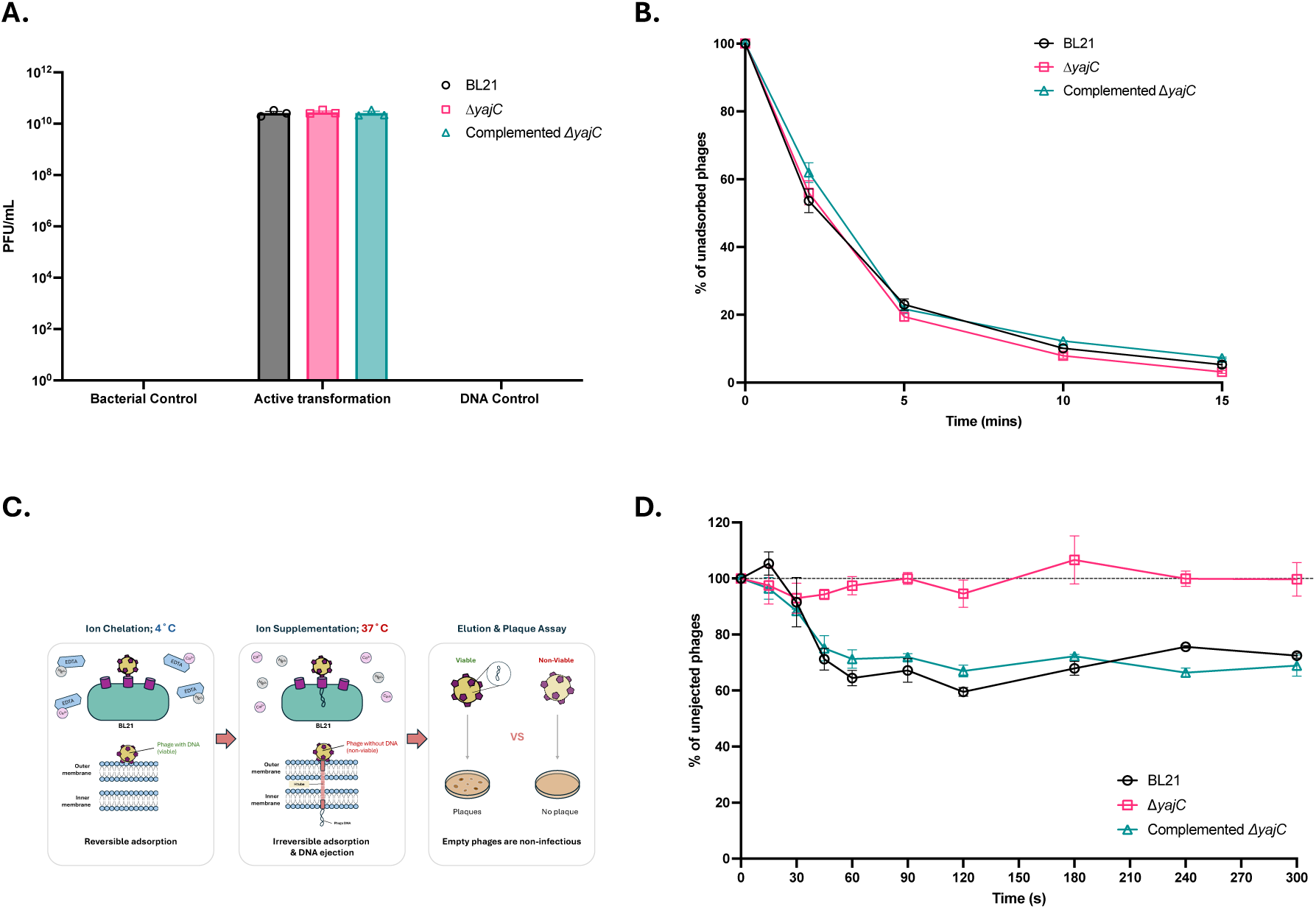
YajC is required for phage genome translocation. **A)** YajC is not required for phage replication and virion assembly. Transformation of phage DNA into Δ*yajC* mutant causes a productive *E. coli* BL21 infection. **B)** Phages adsorb equally well to the wild-type *E. coli* BL21, the *E. coli* BL21 Δ*yajC* mutant and that mutant complemented with a plasmid-borne copy of YajC. **C)** The phage genome ejection assay. Phages were first allowed to adsorb onto the host reversibly at 4°C and under ion-chelated conditions. Divalent cations were then supplemented, and the temperature was raised to 37°C to promote irreversible binding and genome ejection. Given that ejected phages are non-viable, they are unable to infect, hence no plaque formation. **D)** Phage genome ejection kinetics over the first five minutes, in *E. coli* BL21, Δ*yajC* and complemented Δ*yajC*, expressed as the percentage of phages that have not ejected their genomes. All data were presented as mean ± standard error.

### LPS disruption reduces fitness and resensitises phage-resistant bacteria to erythromycin

Next, we determined whether the phage-resistant bacteria paid a growth penalty compared to the parental strain *E. coli* BL21. We found that mutations in LPS genes caused a reduced growth rate (One Way ANOVA; F = 4.79, p = 4 x 10^−4^) and carrying capacity (One Way ANOVA; F = 20.97, *p* = 1 x 10^−4^; Fig. S5) of the phage-resistant bacteria relative to the ancestor, while the *yajC* mutant remained unaffected. We then tested whether phage-resistant bacteria with LPS mutations were more susceptible to antibiotics. We focused on erythromycin, a macrolide antibiotic (*34*) that is primarily effective against Gram-positive bacteria with limited activity against Gram-negative bacteria, due to LPS, which limits the entry of macrolides (*21, 34, 35*). We found that phage-resistant bacteria with LPS mutations were more sensitive to erythromycin, with at least a 4 to 128-fold reduction in resistance, and susceptibility to the antibiotic increased with the severity of LPS disruption (Fig.3A, Table S8), consistent with previous results (*18, 36, 37*). In contrast, the erythromycin susceptibility of the phage-resistant *yajC* mutants was similar to that of the ancestor *E. coli* BL21 (Table S8). In other words, the trade-off between phage and erythromycin resistance in *E. coli* BL21 is only observed for *E. coli* with mutations that disrupt LPS synthesis, but not phage-resistant *E. coli* mutants with mutations in *yajC*.

**Figure 3:**
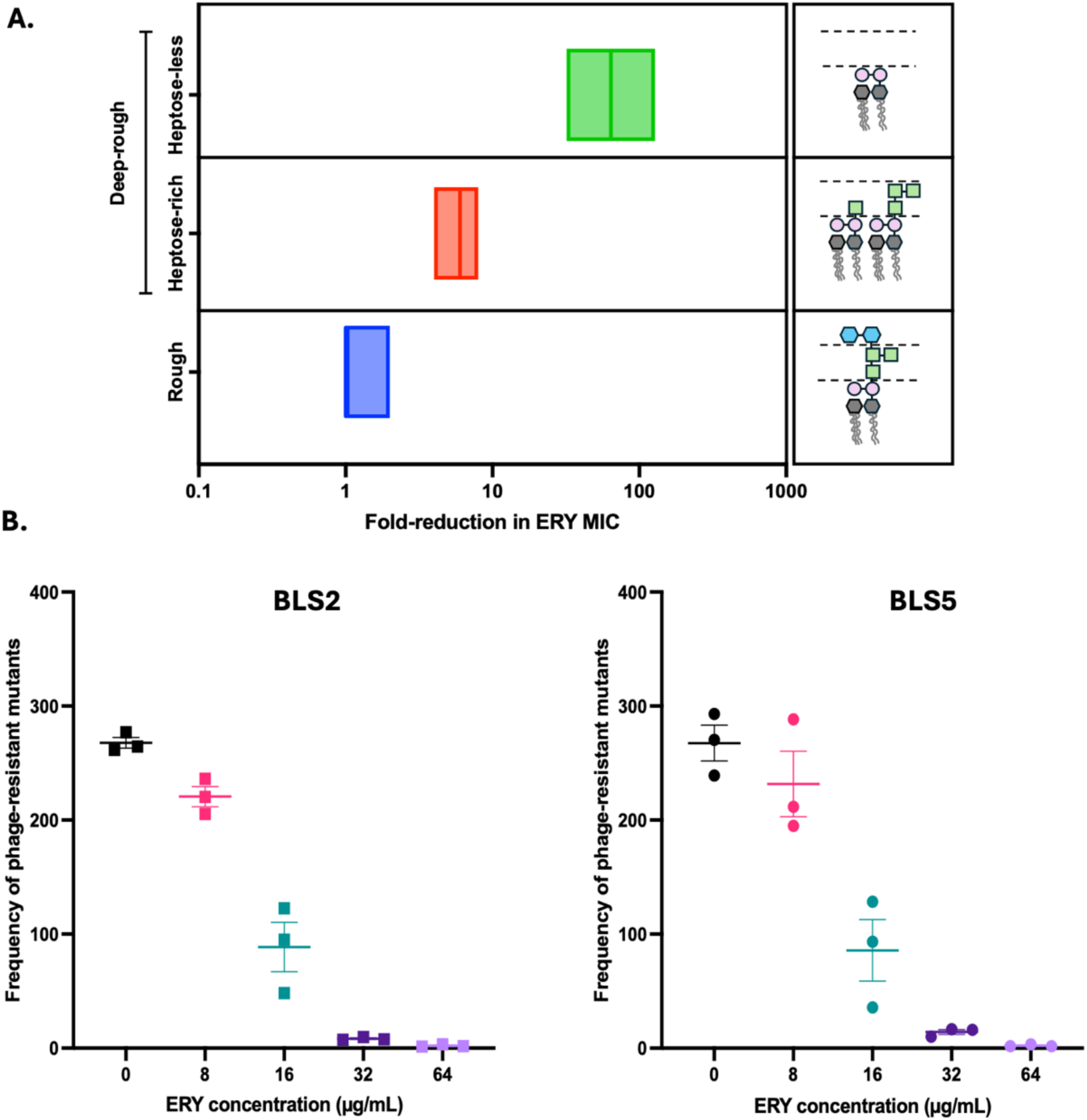
Erythromycin-phage combinations reduce the rate of phage resistance evolution. **A)** Degree of erythromycin susceptibility for the three classes of phage-resistant LPS mutants and **B)** the number of phage-resistant colonies that arise when phage and antibiotic are applied by drop method to a standardised *E. coli* BL21 population (methods).

We also tested whether combining the macrolide antibiotic erythromycin with phage would reduce the evolution of phage resistance. In agreement with previous studies (*38*), we observed that as erythromycin concentration increases, the frequency of phage-resistant bacteria recovered from the population decreases (Fig. 3B). However, the utility of a phage-erythromycin combination would be limited due to the evolution of high-fitness phage-resistant *yajC* mutants that are also resistant to erythromycin (Table S8, Fig. S5).

To address this issue, we employed phage training, a method for directing the evolution of phage towards desirable traits such as expanded host range, high infectivity, or slowed resistance evolution (*5, 11, 39, 40*). We aimed to evolve phages that target the inner core of the LPS and do not target YajC. We reasoned that the only way bacteria could evolve resistance to a phage with these characteristics was by mutations that caused the loss of most LPS, thereby causing sensitivity to erythromycin. Our goal then was to direct the evolution of phage, which could, in turn, steer bacterial evolution towards greater antibiotic sensitivity. To test this idea, we trained the phages BLS2 and BLS5 using *E. coli* BL21, the original host of isolation, in co-culture with one of three *E. coli* strains that the phage could not infect (Table S9). In this co-culture approach, the ancestral host facilitates the maintenance of a large population, while the second, less, or inaccessible host represents an ecological opportunity for the rare mutant to establish a productive infection (*15*) (Fig. 4A). Altogether, we had three treatments. The “K12 treatment” involved training phage to infect an *E. coli* strain, JW5028, with a starkly different LPS structure from the BL21 ancestor (Fig. S6), as a control for the specificity of phage training. Our second treatment was built on the knowledge gained from our characterisation of YajC (Fig. S6). Since mutations in *yajC* confer escape from both phage and antibiotic, with no growth penalty, we aimed to evolve phages that did not require *yajC* for infection and thereby would not drive the evolution of *yajC*-mutant, resistant hosts. Finally, the “LPS treatment” involved training phage to infect *E. coli* BL21 with partial LPS (Fig. S6). After five cycles of evolution, all populations derived from phage BLS2 and phage BLS5 evolved to infect the other less permissive *E. coli* strains (Fig. 4B – C), Fig. S7A – B). It took significantly longer for phage to evolve the capacity to infect the phage-resistant *yajC* mutant (For BLS2, *E. coli* JW5028 = 1.63 ± 0.26 days, *ΔyajC* = 2.63 ± 0.26 days; for BLS5, *E. coli* JW5028 = 1.63 ± 0.38 days, *ΔyajC* = 2.25 ± 0.31 days, Fig. 4D). All evolved phage populations retained their abilities to infect the *E. coli* BL21 ancestor (Fig. S7C – D).

**Figure 4:**
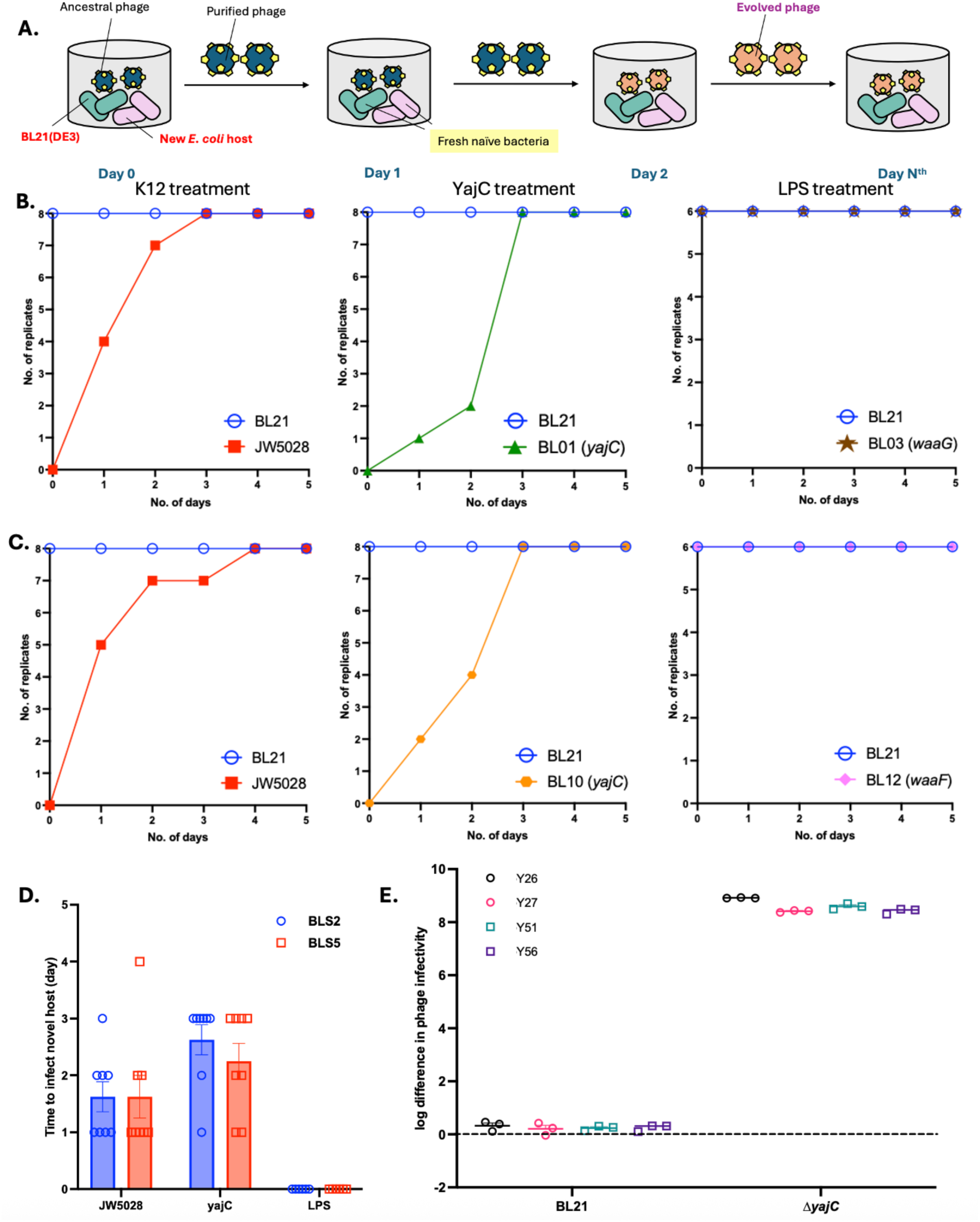
Directed phage training. **A)** Phages were trained in the presence of *E. coli* BL21 (green) and a non-permissive *E. coli* (pink). Bacteria were discarded after each passage, but phages were purified and transferred to another culture of naïve hosts. **B)** Number of BLS2 and **C)** BLS5 phage population replicates that could infect the less permissive *E. coli* host over the phage training period, and **D)** the average time (in days) required for the phage to adapt to new hosts. The mutated gene of the less permissive host is indicated in parentheses. **E)** Infectivity of trained phage against Δ*yajC*. The changes in phage titer before and after training were expressed in log differences; the key indicates the replicate populations. All data were presented as mean ± standard error.

### Directed phage training slows the evolution of phage resistance and steers *E. coli* BL21 towards erythromycin sensitivity

We evaluated the ability of the trained phage populations to suppress *E. coli* BL21 growth without driving the evolution of phage resistance (Table S9), finding significant variation across the three phage training approaches (evolved BLS2: One-way ANOVA, F = 50.61, *p* < 1.0 ×10^−4^; evolved BLS5: One-way ANOVA, F = 11.03, *p* = 1.0 ×10^−4^) (Fig. 5A–B, Supplementary Data 5A – B). All the phage populations from the LPS treatment that we tested had significantly improved capacity to suppress phage resistance evolution compared to the ancestor (Fig. 5A–B, *p* < 0.05), while 1 out of 4 tested phage populations from the YajC treatment were able to suppress resistance evolution compared to the ancestor (Fig. 5**A**–**B**, Supplementary Data 5A–B).

**Figure 5:**
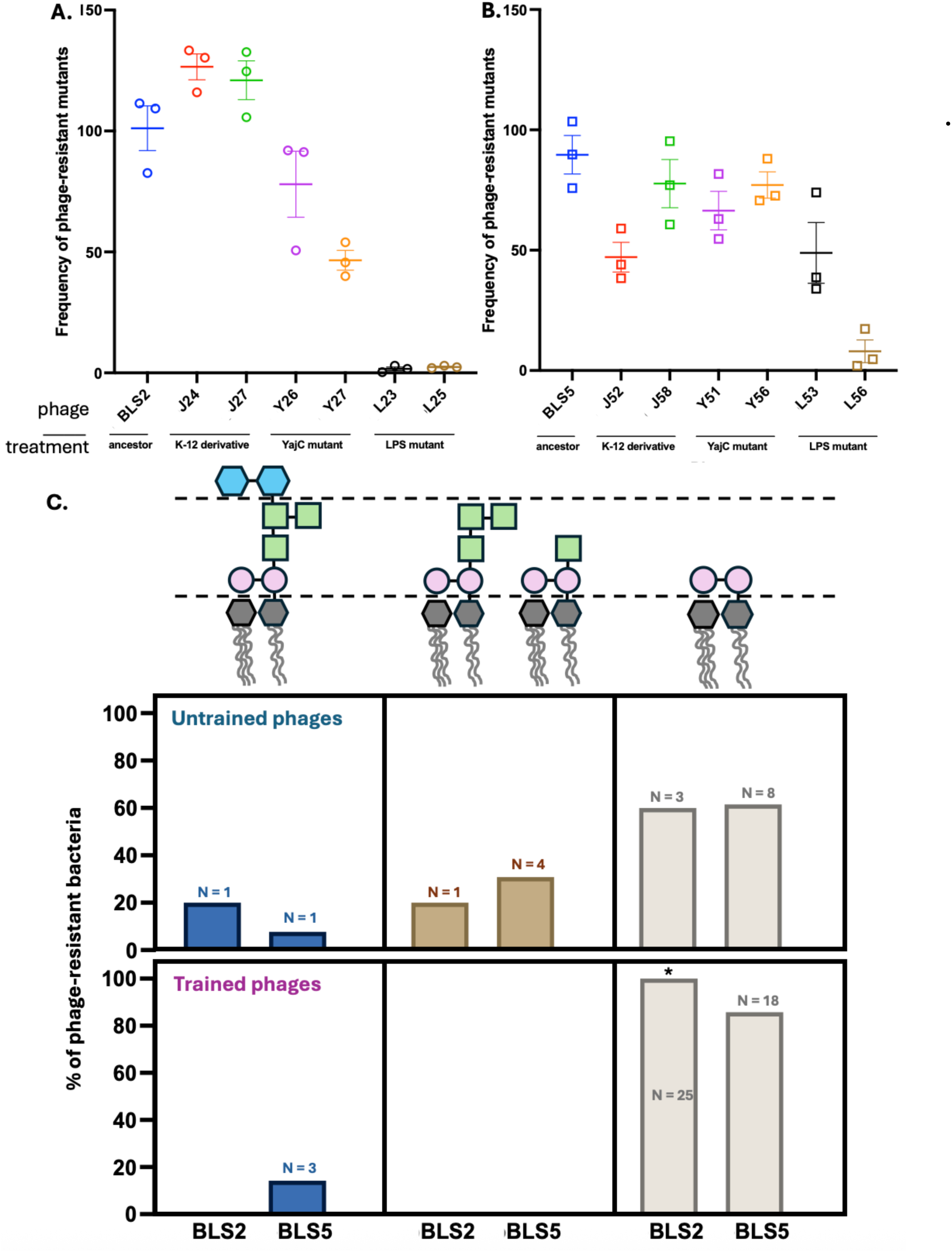
Evolution of resistance to trained phages. **A**) The frequency of phage-resistant mutants that evolved against the trained phage from BLS2 and **B**) BLS5. Statistical significance was determined using One-way ANOVA and Dunnett’s post-hoc test to compare each trained phage to the ancestral phage. Y27 (p = 7 x 10^−4^), L23 (p < 1 x 10^−4^), L25 (p < 1 x 10^−4^), J52 (p = 1.3 x 10^−2^), L53 (p = 1.7 x 10^−2^) and, L56 (p < 1 x 10^−4^). **C)** Distribution of LPS phenotypes of the phage-resistant bacteria upon the evolution of resistance to untrained (top) and trained phages (bottom). The increase in the proportion of heptose-rich deep-rough LPS (grey) in the trained BLS2 population was determined using a one-sided Fisher’s exact test, * P<0.05. N denotes the number of sequenced samples.

Next, we isolated a total of 48, independently evolved, phage-resistant bacteria and performed whole-genome sequencing to identify the genetic causes of resistance to the trained phage (Table S10). We found that all phage-resistant mutants evolved after treatment with the trained BLS2 phage had LPS mutations that result in sensitivity to erythromycin (Fig. 4C, Tables S11 – 12, Supplementary Data 5C). For the phage-resistant mutants evolved after treatment with the BLS5-trained phage, only three 3/21 mutants had antibiotic-resistant genotypes (Fig. 4C, Tables S11 – 12, Supplementary Data 5C). These three antibiotic-resistant mutants evolved in the LPS treatment, while phages trained on the YajC mutant were inevitably antibiotic sensitive.

Overall, this shows that the outcome of the phage training approach depends on two factors. First, was the choice of phage. While most trained phages derived from both ancestor phages improved their capacity to eradicate *E. coli* BL21, phage BLS2 evolved to always drive the evolution of antibiotic sensitivity. It is important to note that after phage training, we re-sequenced the founding phage populations and found that a proportion of the BLS5 population has a C2536T substitution, effectively, the BLS2 genotype; however, we do not think this changes the conclusions of the study. Second, the genotypes used in the training regime led to important differences. The “LPS treatment” populations evolved to target their focal host much more quickly than the YajC treatment populations (Fig. 4D) and suppressed the evolution of phage resistance in the bacterial host more strongly (Fig. 5A – B). However, the YajC treatment populations evolved the capacity to infect *E. coli* hosts that do not have the YajC protein (Fig. 4E) – so that the only path to phage resistance for a host population treated with those trained phage were mutations in LPS that cause antibiotic sensitivity.

## Conclusion

Selection strategies that exploit trade-offs to bias the outcomes of bacterial evolution are are hampered by bacteria evolving down pathways that avoid the trade-off (*3, 9, 10*). Here, we describe a new strategy – directing the evolution of phage to steer bacterial evolution down the genetic route to antibiotic sensitivity. This approach could revitalise the phage steering approaches developed by others (*2, 3, 18*) and, since LPS is a target for many phage that target Gram negative bacteria such as *E. coli* (*9, 30*)*, Klebsiella* (*14, 31*), *Salmonella* (*29*), and *Pseudomonas* (*28*), this approach has the potential to be applied to a wider range of phage.

## Data availability

Whole-genome sequences generated in this study have been deposited in GenBank under the Bioproject ID: PRJNA1023613.

## Author contributions

SKKL carried out experiments, data analysis and visualisation. SKKL and CB carried out DNA sequencing and analysis. SKKL and MJM designed the study and interpreted the data. SKKL, HST and MJM wrote the paper.

## Methods

### Strains and culture conditions

All the *E. coli* strains used in this study were grown overnight at 37°C in LB medium with shaking at 230 rpm. The overnight cultures were sub-cultured for 2 hours after a 2-fold dilution before experimentation unless otherwise stated. *E. coli* JW5028 is a derivative of *E. coli* K-12 BW25113, in which a pseudogene has been replaced by a kanamycin resistance cassette (*41*). *E. coli* BW25113 is the parent strain of the Keio collection of single-gene knockout mutants, making it a suitable reference for functional genomics and genetic studies. *E. coli* JW5028, with a K-12 background, was expected to have a more complex LPS than *E. coli* BL21 (DE3), referred to as *E. coli* BL21 thereafter (Fig. S6). This LPS phenotype likely explains its resistance to phages isolated in this study. All the phage-resistant bacteria were generated in this study. Phages were grown in LB medium supplemented with 10 mM CaCl_2_ and 10 mM MgSO_4_ (referred to as supplemented LB broth), and 0.6% to 0.75% soft LB agar was used for the double agar overlay assay.

### Phage isolation and purification

Aliquots of four different wastewater samples were mixed with *E. coli* BL21 and supplemented with 10X LB medium. Following overnight incubation at 37°C with shaking at 230 rpm, the resulting lysate was purified according to the Phage-on-Tap protocol (*42*) and stored at 4°C. Phage titer (PFU/mL) was quantified using the double agar overlay assay.

### Phage killing assay

Bacterial cultures were adjusted to approximately 1×10^6^ CFU/mL and infected with phages at different multiplicities of infection (MOI) ranging from 1×10^−5^ to 100. Then, 132 µL of each culture was transferred to randomly assigned wells of a 96-well plate. The optical density at 600 nm (OD 600nm) was measured at 37°C every 10 minutes using a microplate reader for 24 hours to track bacterial growth.

### Isolation of phage-resistant bacteria

Phage-resistant bacteria were generated using the double agar overlay approach with 100 µL of phage lysate (≥ 10^8^ PFU/mL). After overnight incubation at 37°C, bacterial colonies that grew were streaked on LB agar. This process was repeated twice for pure colonies isolation (*3*). The phage susceptibility of the phage-resistant bacteria was assessed using the double agar overlay assay.

### Genomic DNA extraction and whole genome sequencing

Bacterial and phage genomic DNA was extracted using the GenElute Bacterial Genomic DNA kit (Sigma-Aldrich) following the manufacturer’s protocol. For phage DNA extraction, additional DNase treatment, isopropanol precipitation, and wash steps were included. Specifically, 5 µL of DNase, 20 µL of RNase, and 50 µL of DNase buffer were added to 1 mL of clarified phage lysate to remove bacterial host DNA and RNA. The mixture was incubated for 2 hours at 37°C, followed by heat inactivation at 75°C for 5 minutes. Then, the lysate was treated with lysis solution and proteinase K at 55°C for 1 hour. Proteinase K was then inactivated at 65°C for 15 minutes. Phage DNA was precipitated using isopropanol, washed four times and purified using a spin column. Genomic DNA for both phages and bacteria were sequenced using the MinION long-read sequencing platform by Oxford Nanopore Technology.

### Bioinformatics analysis

The raw sequencing reads were base-called and demultiplexed using Dorado, retaining bases with a q-score > 10. Next, barcodes and adaptors were removed using Porechop. For variant calling, the Breseq package (*43*) was used to compare the genomes of the phage-resistant bacteria to the parental strain *E. coli* BL21(DE3) as a reference. All the mutations identified in this study have a probability score of 100% (Tables S3 – 4). A total of 19 phage-resistant bacteria genomes were sequenced. No mutation was predicted for one of the BLS2-resistant bacteria. This strain was listed in the unclassified category.

For the phage DNA, the trimmed reads were assembled using Flye (*44*) and polished using Medaka for error correction. Then, the phage contigs were rearranged using the best hit from the Blastn (*45*) output as reference, and the average nucleotide identity (ANI) was computed using FastANI (*46*) for taxonomic classification. Phages were assigned to the same genus with an ANI score > 95%. Gene calling was performed using the annotation tools Prokka (*47*) and Pharokka (*48*). Phage genes were also annotated using Glimmer3, MetaGene Annotator, and Get ORFs, and their functions were predicted using Blastp and InterProScan found in the Phage Galaxy Pipeline (*49*). The draft genomes for BLS2 and BLS5 were visualised using Proksee. However, we acknowledge that the phage genomes may be incomplete as we failed to obtain circularised genomes.

### Gene knockout and complementation

All the primers used to construct the knockout mutant and complementation plasmid were listed in Table S5. The Δ*yajC* knockout mutant was constructed using the λ-red recombination system. Homology fragments (300 bp upstream and downstream of *yajC*) and a chloramphenicol resistance cassette were amplified and combined using overlap extension PCR to generate the Δ*yajC* homology template. *E. coli* BL21 electrocompetent cells were prepared. Briefly, bacteria carrying the λ-red recombination system on the pGETrec plasmid were diluted 1:100 and grown at 37°C with shaking at 200 rpm to an OD 600 nm of 1.0. Then, L-arabinose at a final concentration of 0.2% (w/v) was used to induce the expression of recombinase at 25°C for an additional 40 minutes.

The induced culture was pelleted by centrifugation (5,000 rpm, 10 minutes, 0°C), and the cells were washed four times with 10% (v/v) ice-cold glycerol. After the final wash, the cells were resuspended in 100 µL of 10% glycerol. Then, 20 µL of the cell suspension was mixed with approximately 600 ng of Δ*yajC* template. The mixture was transferred to a sterile 0.2 cm electroporation cuvette and pulsed at 2.5 kV, 200 ohms, and 25 µF. Super-optimal broth pre-warmed to 37°C was added immediately, and the cells were recovered for one hour at 37°C. The recovered cells were pelleted, resuspended in 100 µL of LB broth, and spread-plated onto LB agar supplemented with 12.5 µg/ml chloramphenicol. Potential Δ*yajC* knockout colonies that grew under chloramphenicol selection were PCR-amplified using primers (Table S5) and NEB Standard Taq polymerase to confirm successful gene deletion. The pGETrec plasmid was then cured from Δ*yajC* mutant.

To complement *yajC* back into Δ*yajC the* pUltra vector, the *yajC* gene and its native promoter were amplified using primers with EcoRI and XbaI sites (Table S5). The amplified vector was treated with DpnI (New England Biolabs). Both vector and insert fragments were then purified using the Monarch PCR & DNA Cleanup Kit and digested with EcoRI and XbaI (Vivantis Technologies). The digested products were ligated using T4 DNA Ligase (New England Biolabs) at a vector-to-insert molar ratio of 1:5. The ligated product was chemically transformed into *E. coli* BL21 competent cells. Transformants were selected on LB agar supplemented with 50 µg/ml of kanamycin and screened using colony PCR. The pUltra-yajC plasmid was extracted and transformed into Δ*yajC*.

### Efficiency of Plating

The efficiency of plating (EOP) assay was used to compare the phage susceptibility of *E. coli* BL21, Δ*yajC*, and the complemented strain. Bacterial cultures were allowed to grow for two hours and were infected with 100 µL of undiluted phage lysate (≈10^9^ PFU/mL). After overnight incubation, the number of plaques was recorded and compared to the plaques formed from the original host *E. coli* BL21.

### Phage adsorption assay

Bacterial cultures were diluted 1:100 in LB and grown at 37°C for two hours. Cells were pelleted by centrifugation (4,000 rpm, 7 mins) and washed twice in adsorption buffer (50 mM Tris, pH 7.4, 100 mM NaCl, 100 mM CaCl_2_, 8 mM MgSO_4_). Bacterial culture and BLS2 lysate were mixed at an MOI of 0.1 in the adsorption buffer. Samples were taken at the designated time point, centrifuged immediately at 4,000g, 4°C for five minutes to pellet the bacterial cells and remove any adsorbed phages. The supernatant was diluted, and the quantity of free phages was determined using the double agar overlay assay.

### Phage DNA replication and virion assembly

Phage DNA was amplified, and approximately 100 ng of the DNA was mixed with 50 µL of chemically competent cells and incubated on ice for 25 minutes. The mixture was heat-shocked at 42°C for 90 seconds and immediately kept on ice for 5 minutes. The cells were recovered in 5 mL supplemented LB for two hours at 37°C to facilitate phage assembly. The culture was centrifuged at 4,000 rpm for 15 minutes, and 4 mL of the supernatant was transferred to 400 µL of the original host of isolation, *E. coli* BL21. The mixture was incubated at 37°C with shaking at 200 rpm for an additional five hours to amplify phage production. Following incubation, the culture was centrifuged at 13,000 g for 15 minutes to recover the phage lysate. The lysate was diluted and plated to check for the presence of phages.

### Phage-mediated cell lysis

The pBAD24 vector with protein E, pBAD24-E, was transformed into *E. coli* BL21, Δ*yajC,* and the complementation strains were maintained in the presence of 100 µg/mL ampicillin. The cultures were diluted 1:100 and grown until the mid-log phase. Then, they were standardised to an OD 600 nm of 0.2 and induced with 0.2% L-arabinose. The bacterial growth was monitored using a microplate reader (Tecan Infinite Pro) at 30-minute intervals until lysis was essentially completed. Bacteria with the pBAD24-hgLA vector served as a negative control.

### Bacterial growth assays

Growth assays for all the phage-resistant bacteria were conducted using a similar protocol to the phage-killing assay. In brief, bacterial cultures were standardised to 1×10^5^ CFU/mL and 132 µL of each culture was transferred to a 96-well plate. Data was analysed using the Growthcurver package in R to determine the growth rate (r) and maximum carrying capacity (K).

### Minimum inhibitory concentration of erythromycin

The minimum inhibitory concentration of erythromycin for *E. coli* BL21 and the phage-resistant mutants (BL01 to BL12) was determined using the micro-broth dilution approach. Bacterial cultures standardised to 1×10^5^ CFU/mL were treated with different concentrations of erythromycin, ranging from 0.5 µg/mL to 256 µg/mL. Then, 132 µL of each culture was transferred to a 96-well plate and incubated overnight at 37°C. The lowest concentration of erythromycin that did not have any visible bacterial growth was considered the minimum inhibitory concentration.

### Phage-erythromycin assay

The combined effects of phage with erythromycin were determined using the spot assay approach. To keep the bacterial population size consistent, each phage-antibiotic combination was spotted onto the same lawn. If this experiment were conducted in liquid broth, our concern was that the antibiotic could have an impact on the population size, which could cause a lower number of phage-resistant bacteria. Erythromycin was added to 10^7^ PFU/mL of phage to achieve a final concentration of 8, 16, 32, and 64 µg/mL. Then, 5 µL of this mixture was spotted on a bacterial lawn of 10^9^ CFU/mL. After overnight incubation, the frequency of phage-resistant bacteria growing in the clearing zone was determined and compared to phage-only and erythromycin-only controls.

### Directed phage evolution

Phage populations were allowed to evolve in three combinations of a two-host culture setup. The original host of isolation, *E. coli* BL21, was always included in all treatments. For the second host, less permissive *E. coli* with different LPS compositions were used. This included JW5028 and phage-resistant mutants Δ*[yajC]*, *waaG*-C102F and *waaF*-R113H. Training using the JW5028 and Δ*[yajC]* was replicated eight times, while that using *waaG*-C102F and *waaF*-R113H was replicated six times.

Bacteria were mixed in a ratio of 1:1, achieving a total cell density of 1×10^8^ CFU/mL. Then phages were added at an MOI of 0.1. After 24 hours of incubation at 37°C, bacteria were discarded, and phage lysates were purified and transferred to fresh naïve bacterial cultures. Naïve bacteria were defined as bacteria not previously exposed to phages. Only phages were allowed to evolve using this experimental design (*15*). The overall process was conducted for 5 days.

After each evolution, 3 µL of each phage lysate was spotted on the corresponding bacterial lawn to determine if there were any changes in the infectivity of the trained phage populations against the original host and the less-permissive *E. coli*. The presence of a clearing zone was counted as positive for infection (*15*).

### Characterisation of evolved phage population G phage-resistant mutants

Two replicates of evolved phage populations from cycle 5 with strong clearing in the spot assay were amplified using the corresponding second host in each setup and purified (Fig. S7A – 7B). The infectivity of these evolved phage populations against the original host *E. coli* BL21 was determined and compared to ancestral phages BLS2 and BLS5. A double agar overlay assay was also performed to compare the infectivity of the phage communities Y26, Y27, Y51, and Y56 against the original host *E. coli* BL21 and Δ*yajC*. Besides, the infectivity of these evolving phage populations was compared to that of the ancestor BLS2 and BLS5 on both *E. coli* BL21 and Δ*yajC*. Given that the ancestor phages were unable to infect Δ*yajC*, a pseudo-count was introduced to deduce the log differences in phage titer before and after training, as a measure of a change in phage infectivity.

Spot assay was used to compare the ability of *E. coli* BL21 to evolve resistance to both untrained and trained phages. Briefly, 5 µL of the phage lysate adjusted to 1×10^6^ PFU/mL was spotted on *E. coli* BL21 lawn standardised to 1×10^8^ CFU/mL. After overnight incubation, the frequency of phage-resistant bacteria that grew in the zone of clearance was determined. Phage-resistant bacteria were isolated and purified from two re-streaked. The phage susceptibility of phage-resistant bacteria was assessed using an inverted spot assay, where 5 µL of bacterial culture was spotted onto a phage lawn. Genomic DNA from a total of 48 phage-resistant bacteria were extracted, sequenced and analysed as mentioned above.

## Supplementary Information

### Supplementary Tables

**Supplementary Table 1:**
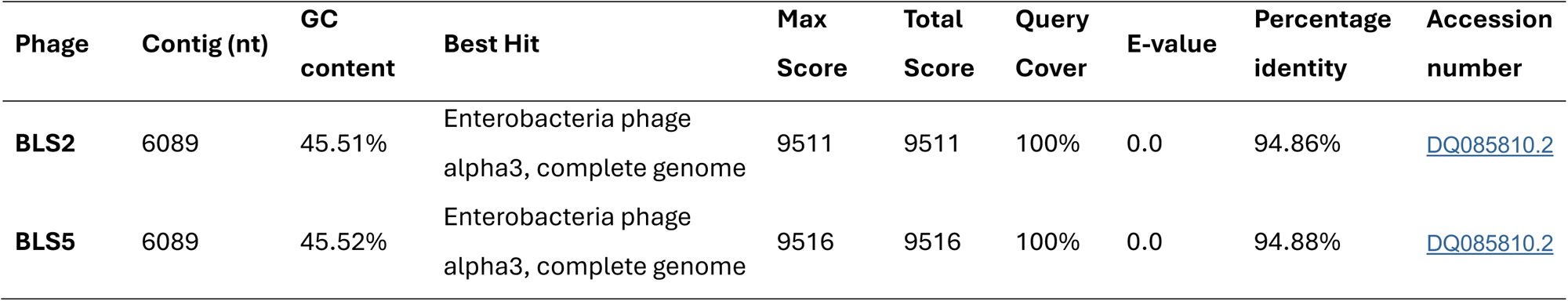
Phage genomic features and sequence alignment from BLAST.

| Phage | Contig (nt) | GC content | Best Hit | Max Score | Total Score | Query Cover | E-value | Percentage identity | Accession number |
| --- | --- | --- | --- | --- | --- | --- | --- | --- | --- |
| **BLS2** | 6089 | 45.51% | Enterobacteria phage alpha3, complete genome | 9511 | 9511 | 100% | 0.0 | 94.86% | DQ085810.2 |
| **BLS5** | 6089 | 45.52% | Enterobacteria phage alpha3, complete genome | 9516 | 9516 | 100% | 0.0 | 94.88% | DQ085810.2 |

**Supplementary Table 2:** Phage genome annotation.

| Gene ID | Type | Gene | Start | End | Length | Strand | Function <sup>b</sup> |
| --- | --- | --- | --- | --- | --- | --- | --- |
| P01 | CDS | *put1*<sup>a</sup> | 66 | 434 | 369 | + | Hypothetical protein |
| P02 | CDS | *put2* | 438 | 527 | 90 | + | Hypothetical protein |
| P03 | CDS | *put3* | 524 | 700 | 177 | + | Hypothetical protein |
| P04 | CDS | *put4* | 418 | 510 | 93 | + | Hypothetical protein |
| P05 | CDS | *put5* | 555 | 671 | 117 | + | Hypothetical protein |
| P06 | CDS | A | 760 | 2244 | 1485 | + | Phage DNA replication |
| P07 | CDS | A* | 1171 | 2244 | 1074 | + | Unknown function |
| P08 | CDS | B | 1806 | 2159 | 354 | + | Internal scaffolding protein |
| P09 | CDS | K | 2159 | 2320 | 162 | + | Unessential; unknown function |
| P10 | CDS | C | 2271 | 2477 | 207 | + | DNA maturation |
| P11 | CDS | D | 2504 | 2956 | 453 | + | External scaffolding protein |
| P12 | CDS | E | 2676 | 2903 | 228 | + | Host cell lysis |
| P13 | CDS | J | 2958 | 3032 | 75 | + | DNA binding protein |
| P14 | CDS | F | 3068 | 4363 | 1296 | + | Major capsid protein |
| P15 | CDS | G | 4398 | 4961 | 564 | + | Major spike protein |
| P16 | CDS | H | 5097 | 6089 | 993 | + | Minor spike protein; DNA pilot protein |

<sup>a</sup> The *put* genes are only specific to alpha3-clade micro-viruses
<sup>b</sup> Functions of the microvirid genes (1)

**Supplementary Table 3:**
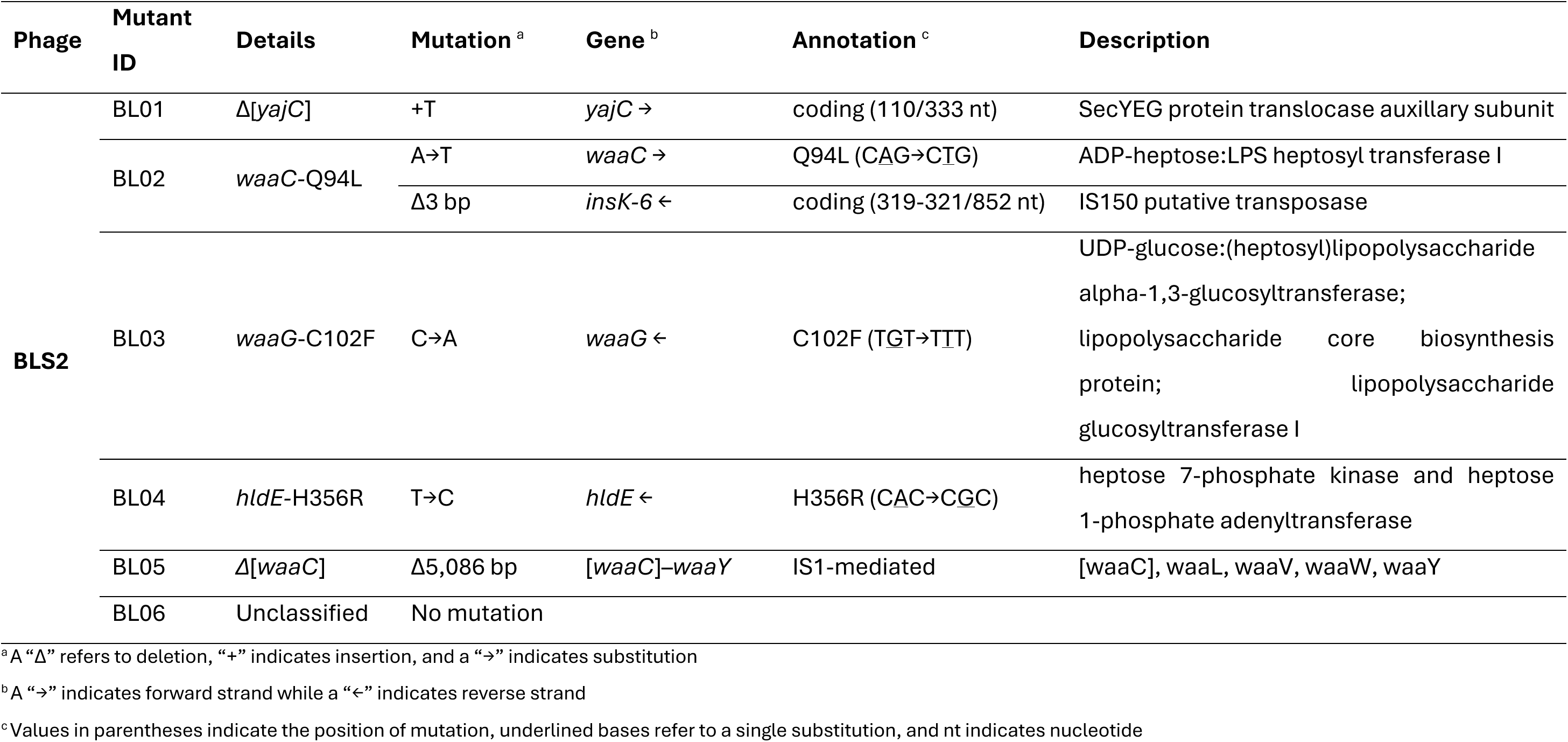
Mutations in phage BLS2-resistant *E. coli* BL21.

**Supplementary Table 4:**
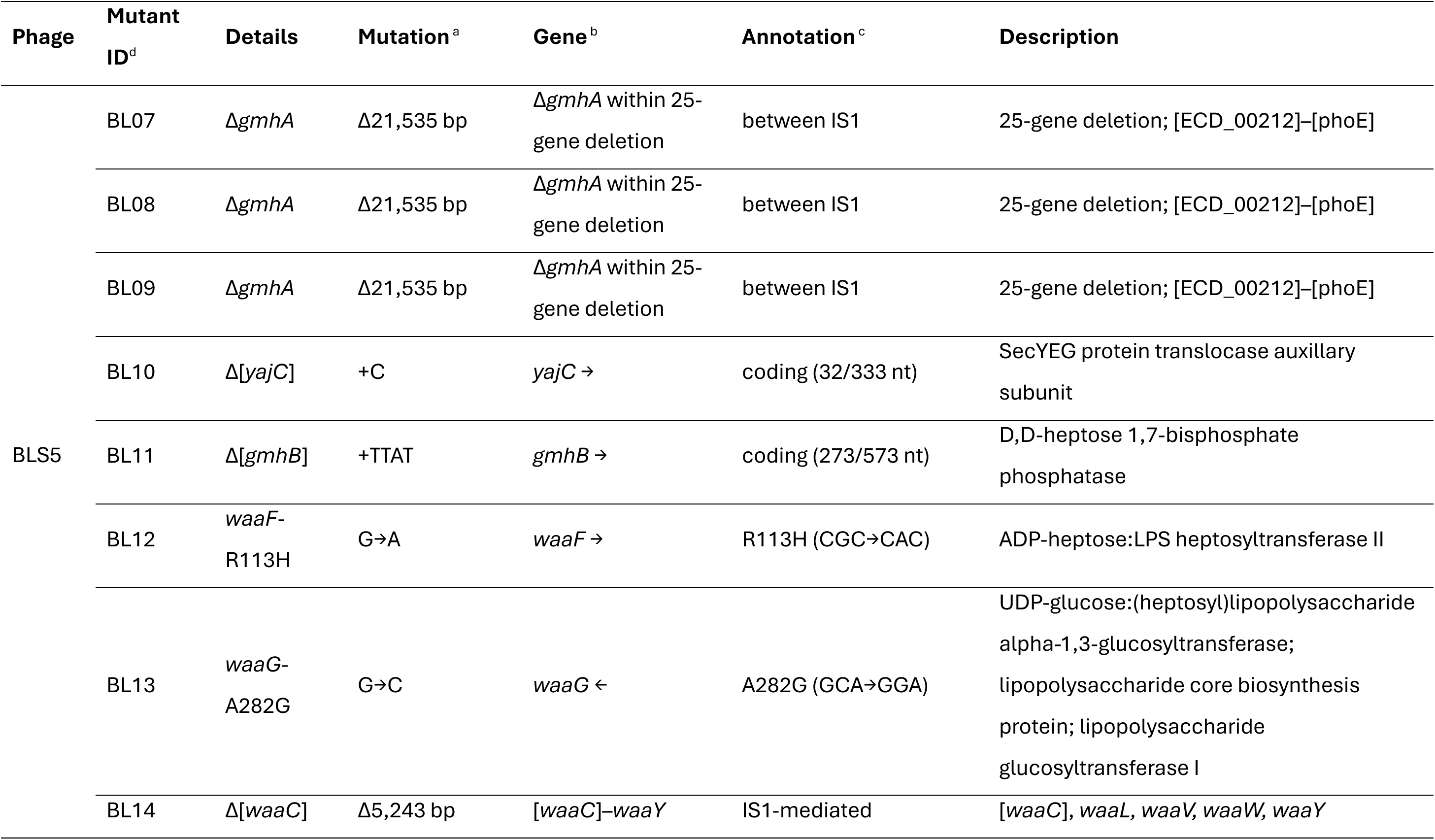
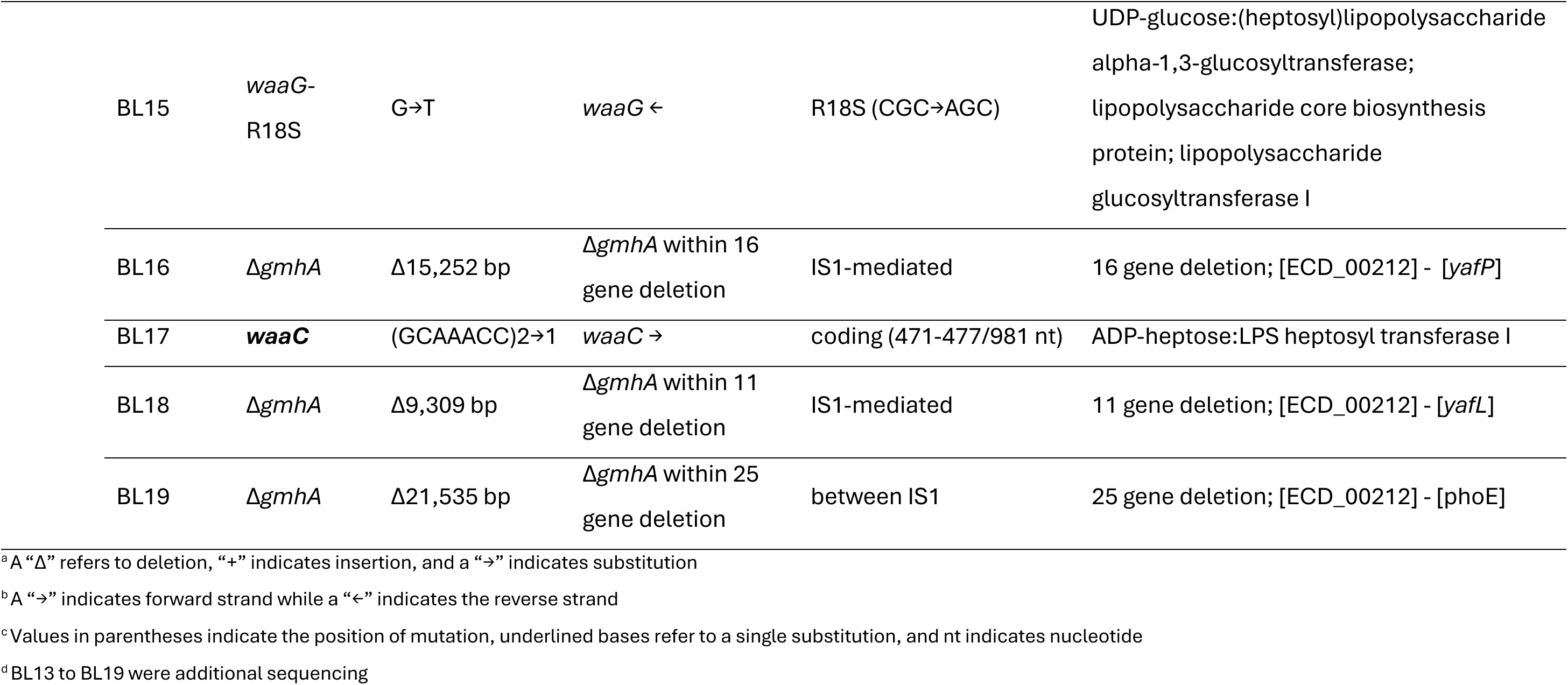
Mutations in BLS5-resistant *E. coli* BL21.

| Phage | Mutant ID<sup>d</sup> | Details | Mutation<sup>a</sup> | Gene<sup>b</sup> | Annotation<sup>c</sup> | Description |
| --- | --- | --- | --- | --- | --- | --- |
| BLS5 | BL07 | $\Delta gmhA$ | $\Delta 21,535$ bp | $\Delta gmhA$ within 25-gene deletion | between IS1 | 25-gene deletion; [ECD_00212]–[phoE] |
| | BL08 | $\Delta gmhA$ | $\Delta 21,535$ bp | $\Delta gmhA$ within 25-gene deletion | between IS1 | 25-gene deletion; [ECD_00212]–[phoE] |
| | BL09 | $\Delta gmhA$ | $\Delta 21,535$ bp | $\Delta gmhA$ within 25-gene deletion | between IS1 | 25-gene deletion; [ECD_00212]–[phoE] |
| | BL10 | $\Delta [yajC]$ | +C | *yajC* → | coding (32/333 nt) | SecYEG protein translocase auxillary subunit |
| | BL11 | $\Delta [gmhB]$ | +TTAT | *gmhB* → | coding (273/573 nt) | D,D-heptose 1,7-bisphosphate phosphatase |
|  | BL12 | *waaF*-R113H | G→A | *waaF* → | R113H (CGC→CAC) | ADP-heptose:LPS heptosyltransferase II |
|  | BL13 | *waaG*-A282G | G→C | *waaG* ← | A282G (GCA→GGA) | UDP-glucose:(heptosyl)lipopolysaccharide alpha-1,3-glucosyltransferase; lipopolysaccharide core biosynthesis protein; lipopolysaccharide glucosyltransferase I |
| | BL14 | $\Delta [waaC]$ | $\Delta 5,243$ bp | $[waaC]$–*waaY* | IS1-mediated | $[waaC]$, *waaL*, *waaV*, *waaW*, *waaY* |

|  |  |  |  |  |  |
| --- | --- | --- | --- | --- | --- |
| BL15 | *waaG*-<br>R18S | G→T | *waaG* ← | R18S (CGC→AGC) | UDP-glucose:(heptosyl)lipopolysaccharide<br>alpha-1,3-glucosyltransferase;<br>lipopolysaccharide core biosynthesis<br>protein; lipopolysaccharide<br>glucosyltransferase I |
| BL16 | $\Delta gmhA$ | $\Delta 15,252$ bp | $\Delta gmhA$ within 16<br>gene deletion | IS1-mediated | 16 gene deletion; [ECD_00212] - [*yafP*] |
| BL17 | ***waaC*** | (GCAAACC)<sup>2</sup>→1 | *waaC* → | coding (471-477/981 nt) | ADP-heptose:LPS heptosyl transferase I |
| BL18 | $\Delta gmhA$ | $\Delta 9,309$ bp | $\Delta gmhA$ within 11<br>gene deletion | IS1-mediated | 11 gene deletion; [ECD_00212] - [*yafL*] |
| BL19 | $\Delta gmhA$ | $\Delta 21,535$ bp | $\Delta gmhA$ within 25<br>gene deletion | between IS1 | 25 gene deletion; [ECD_00212] - [*phoE*] |

<sup>a</sup> A “Δ” refers to deletion, “+” indicates insertion, and a “→” indicates substitution
<sup>b</sup> A “→” indicates forward strand while a “←” indicates the reverse strand
<sup>c</sup> Values in parentheses indicate the position of mutation, underlined bases refer to a single substitution, and nt indicates nucleotide
<sup>d</sup> BL13 to BL19 were additional sequencing

**Supplementary Table 5:**
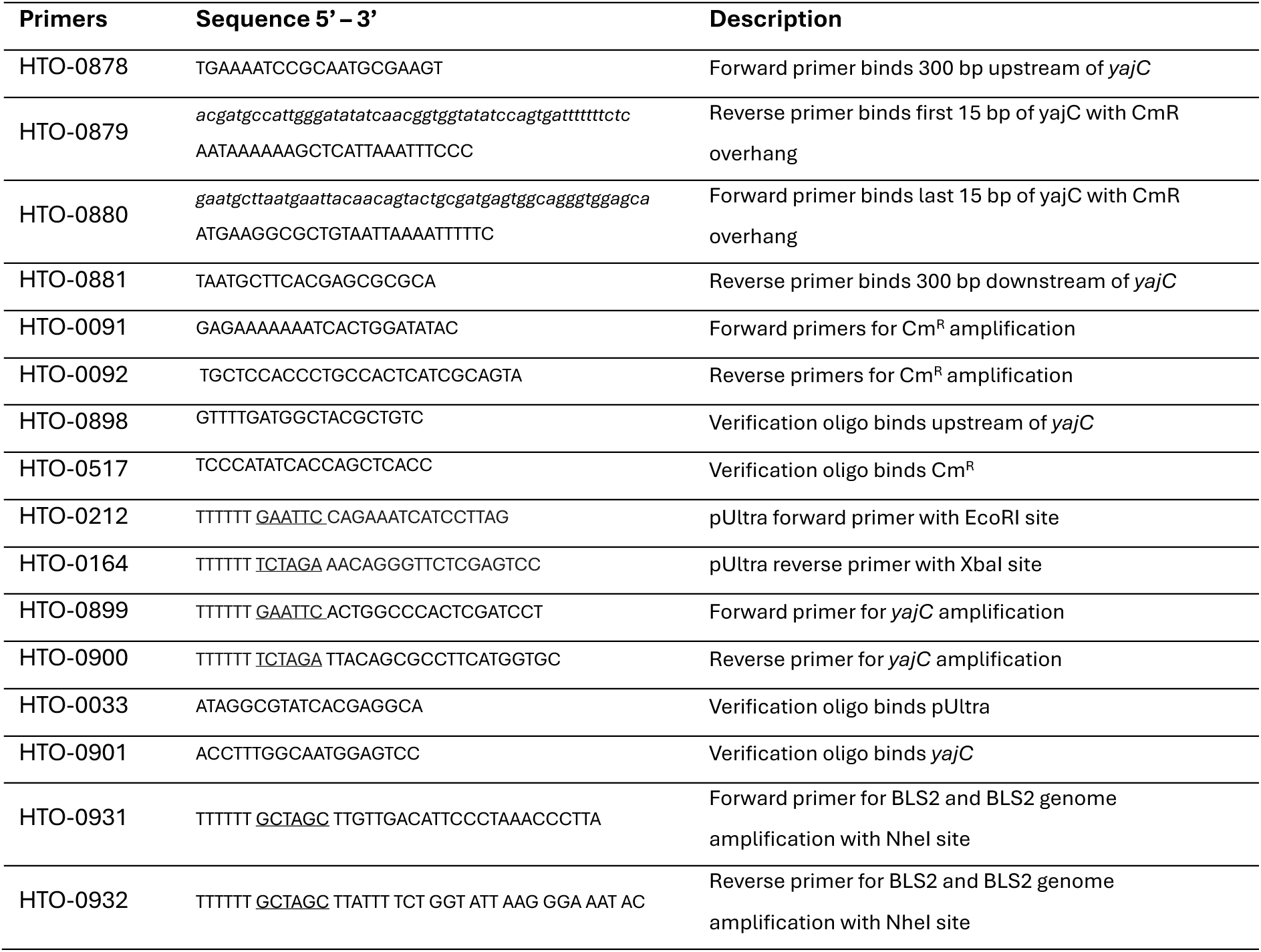
Primers used in this study with the restriction site underlined.

**Supplementary Table 6:** BLS2 infectivity on *E. coli* BL21, Δ*yajC*, and complemented Δ*yajC*.

| Bacterial strain | Phage infectivity (x10<sup>9</sup> PFU/mL) |  |  |  |
| --- | --- | --- | --- | --- |
|  | Rep 1 | Rep 2 | Rep 3 | Mean <sup>a</sup> |
| *E. coli* BL21 | 1.66 | 0.60 | 0.79 | 1.02 ± 0.33 |
| $\Delta yajC$ | 0.00 | 0.00 | 0.00 | 0.00 ± 0.00 |
| Complemented $\Delta yajC$ | 1.02 | 0.66 | 0.75 | 0.81 ± 0.12 |

<sup>a</sup> Data was expressed as mean ± standard error

**Supplementary Table 7:** BLS5 infectivity on *E. coli* BL21, Δ*yajC*, and complemented Δ*yajC*.

| Bacterial strain | Phage infectivity (x10<sup>9</sup> PFU/mL) |  |  |  |
| --- | --- | --- | --- | --- |
|  | Rep 1 | Rep 2 | Rep 3 | Mean <sup>a</sup> |
| *E. coli* BL21 | 1.69 | 1.60 | 2.43 | 1.91 ± 0.27 |
| $\Delta yajC$ | 0.00 | 0.00 | 0.00 | 0.00 ± 0.00 |
| Complemented $\Delta yajC$ | 1.66 | 1.71 | 2.23 | 1.87 ± 0.18 |

<sup>a</sup> Data was expressed as mean ± standard error

**Supplementary Table 8:** Raw data of the minimum inhibitory concentration (MIC) for erythromycin. MIC was determined using the microbroth dilution approach.

| *E. coli* strains | Erythromycin MIC (µg/mL) |  |  |
| --- | --- | --- | --- |
|  | Replicate 1 | Replicate 2 | Replicate 3 |
| BL21WT | 128 | 128 | 128 |
| **BLS2 phage-resistant mutant** |  |  |  |
| $\Delta[yajC]$ | 64 | 64 | 128 |
| *waaC*-Q94L | 2 | 2 | 2 |
| *waaG*-C102F | 32 | 32 | 32 |
| *hldE*-H356R | 1 | 1 | 1 |
| $\Delta[waaC]$ | 2 | 2 | 2 |
| Unclassified <sup>a</sup> | 8 | 8 | 8 |
| **BLS5 phage-resistant mutant** |  |  |  |
| $\Delta[gmhA]$ | 2 | 2 | 2 |
| $\Delta[gmhA]$ | 2 | 2 | 2 |
| $\Delta[gmhA]$ | 2 | 2 | 2 |
| $\Delta[yajC]$ | 128 | 128 | 128 |
| $\Delta[gmhB]$ | 4 | 4 | 4 |
| *waaF*-R113H | 16 | 16 | 16 |

<sup>a</sup> Phage-resistant bacteria with no mutations identified from Breseq

**Supplementary Table 9:** Evolved phage populations selected for downstream analysis.

| Phage | Second *E. coli* host | Evolved phage population <sup>b</sup> |
| --- | --- | --- |
| BLS2 | *E. coli* JW5028 | J24, J27 |
| | $\Delta[yajC]$ | Y26, Y27 |
|  | *waaG*-C102F <sup>a</sup> | L23, L25 |
| BLS5 | *E. coli* JW5028 | J52, J58 |
| | $\Delta[yajC]$ | Y51, Y56 |
|  | *waaF*-R113H <sup>a</sup> | L53, L56 |

<sup>a</sup> Both *waaG*-C102F and *waaF*-R113H were phage-resistant bacteria with LPS-related mutation. For simplicity in naming the evolved phage populations, they were collectively known as LPS mutants.
<sup>b</sup> Phages were named using the following convention: **J**, **Y**, **L** = second host in phage training (*JW5028*, *yajC*, *LPS* mutants); 1<sup>st</sup> number (**2** or **5**) = ancestral phages (BLS**2** or BLS**5**); 2<sup>nd</sup> number = replicate number.

**Supplementary Table 10:**
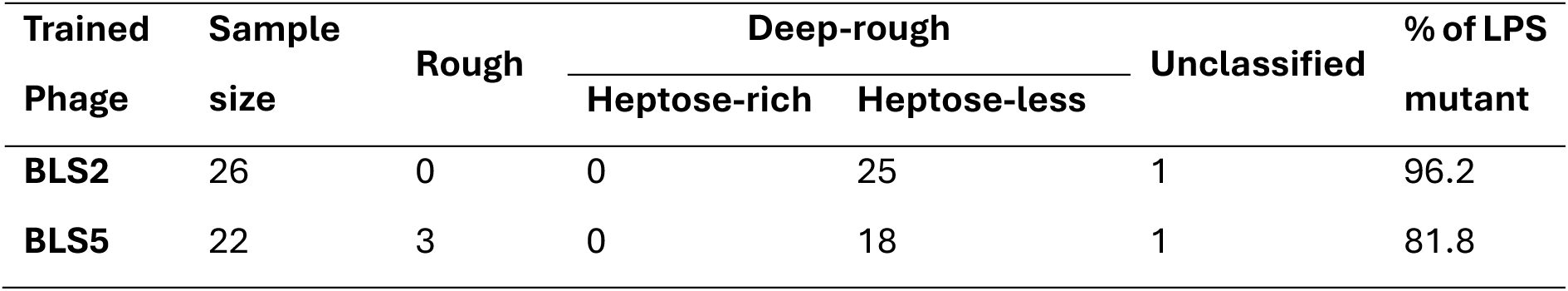
LPS distribution of bacteria resistant to evolutionary trained phages.

**Supplementary Table 11:**
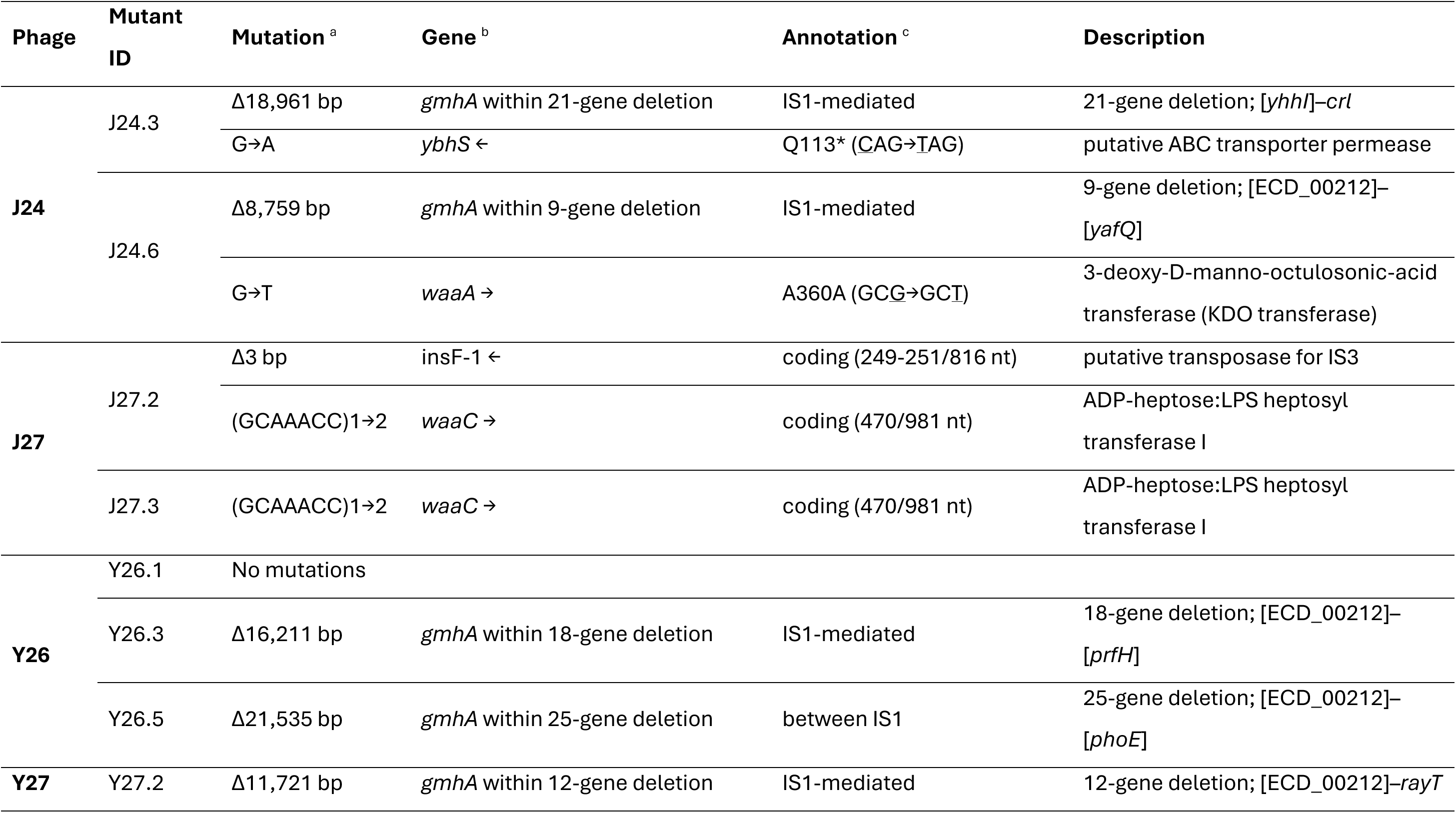
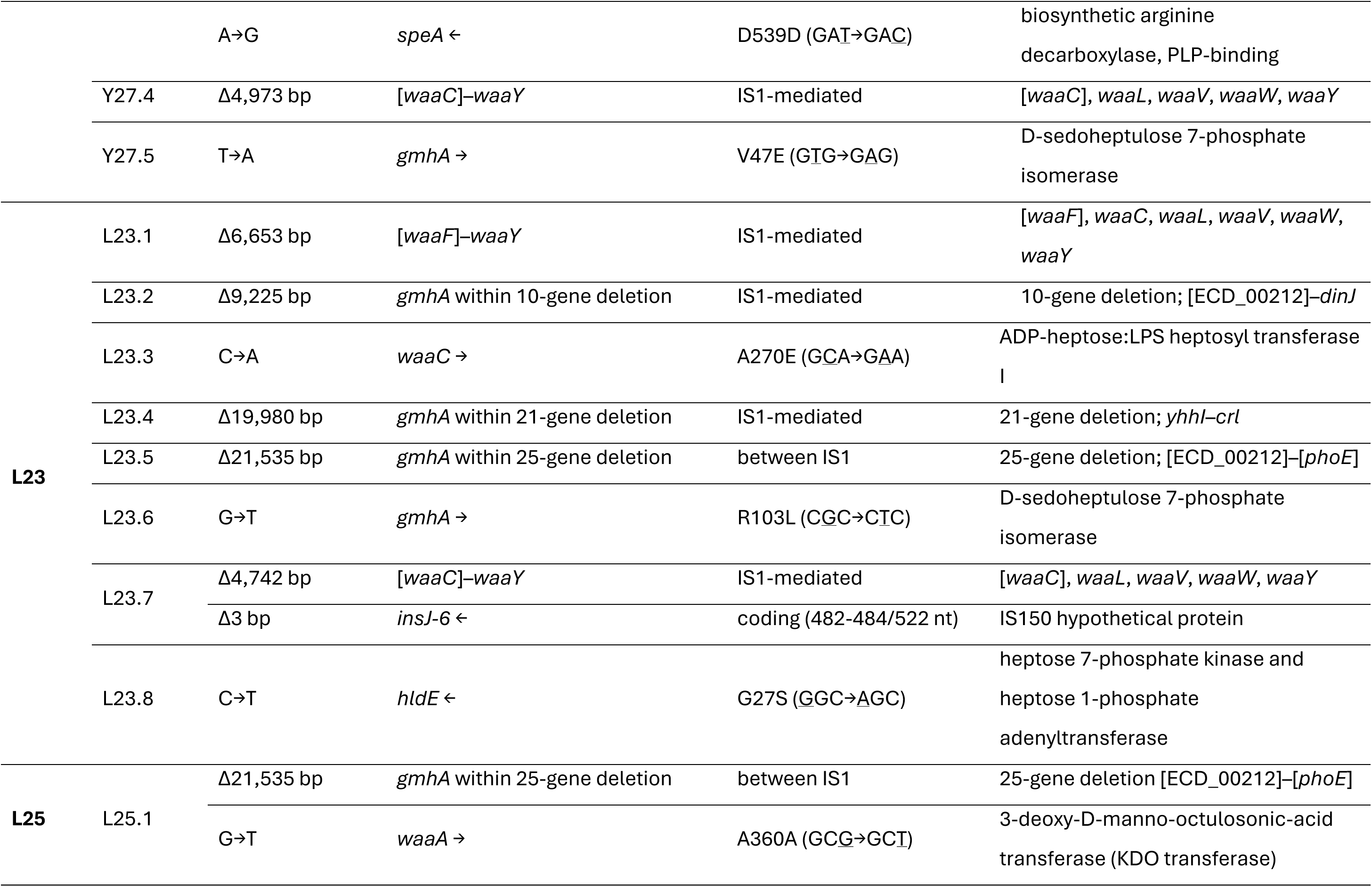
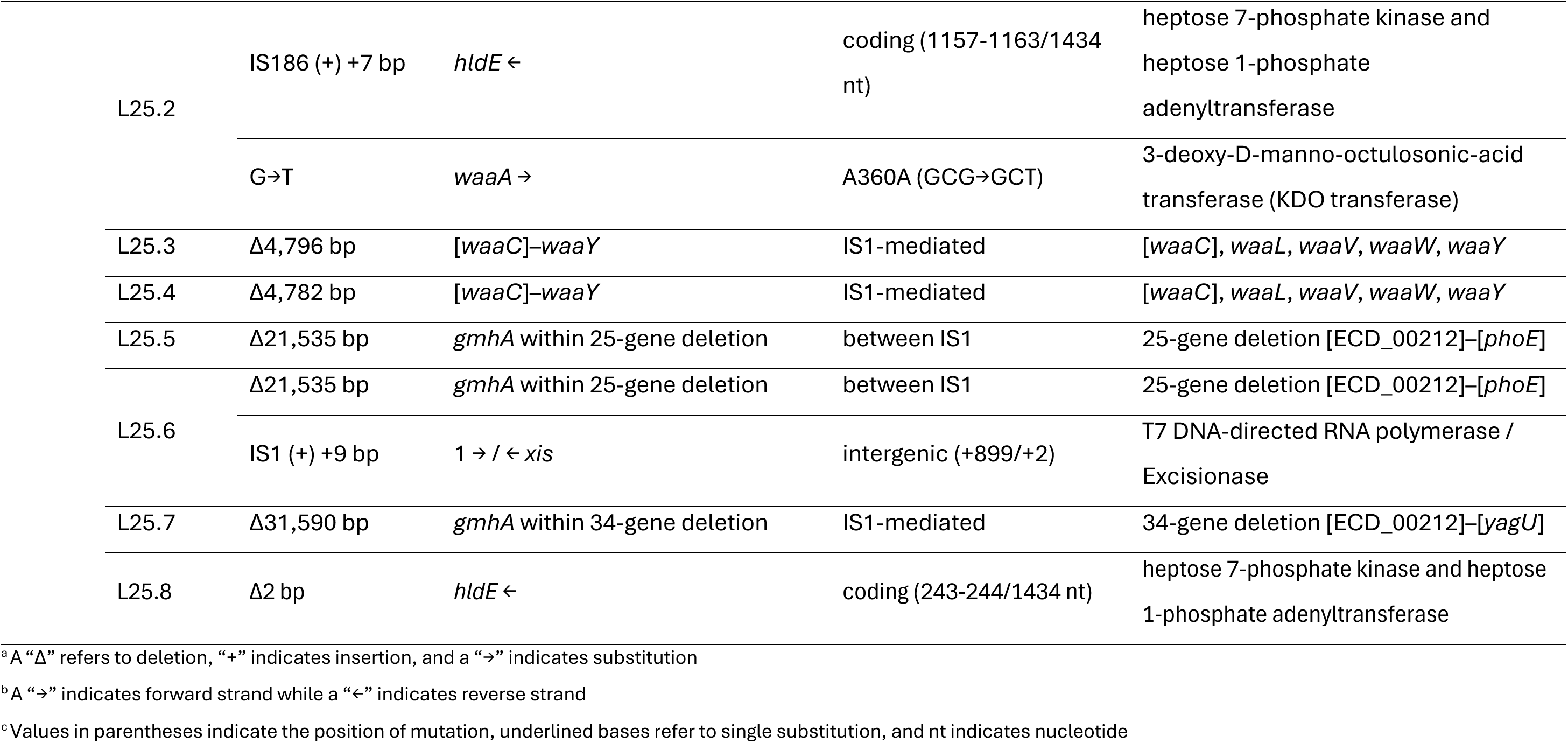
Phage-resistant mutations against evolutionary trained BLS2.

**Supplementary Table 12:**
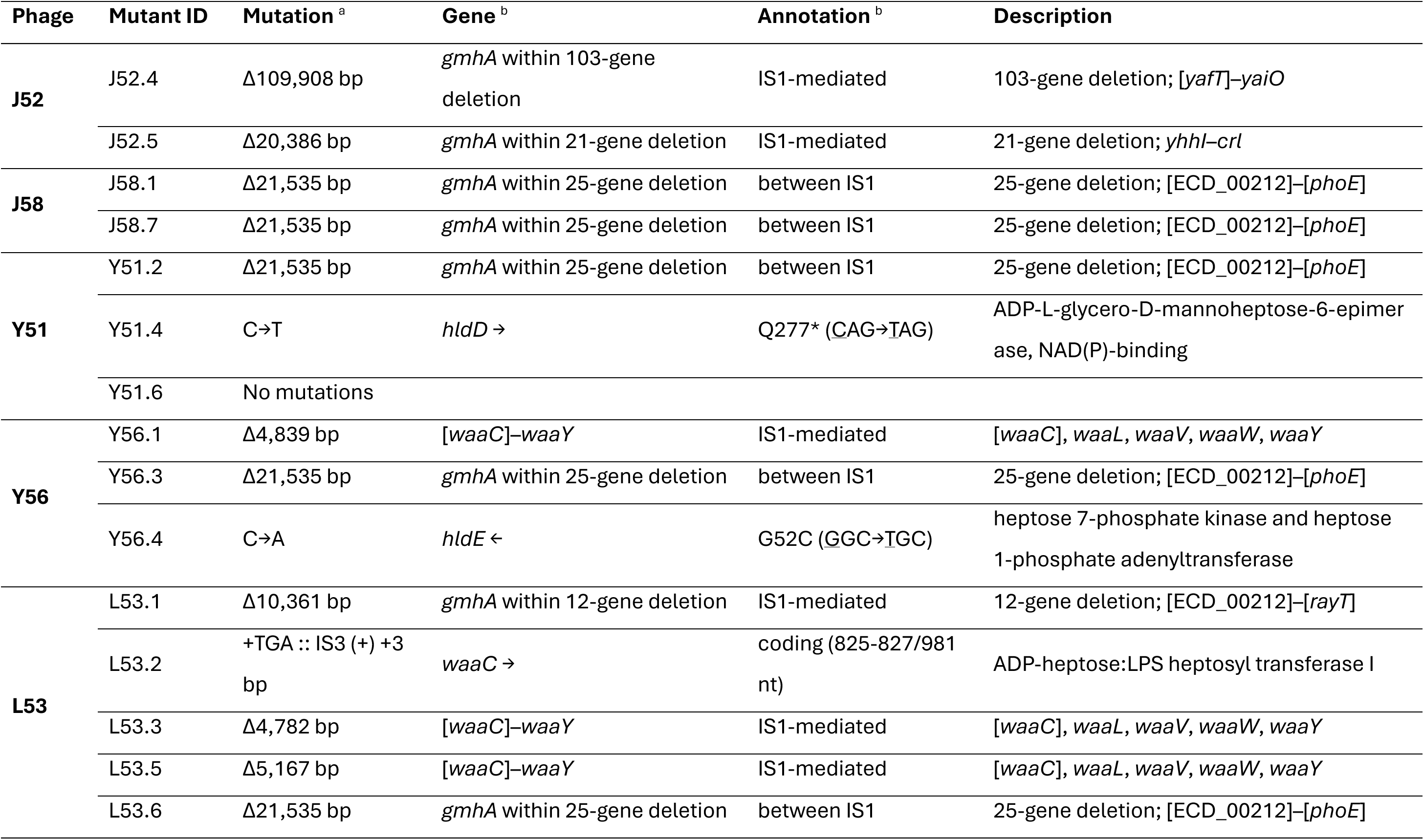
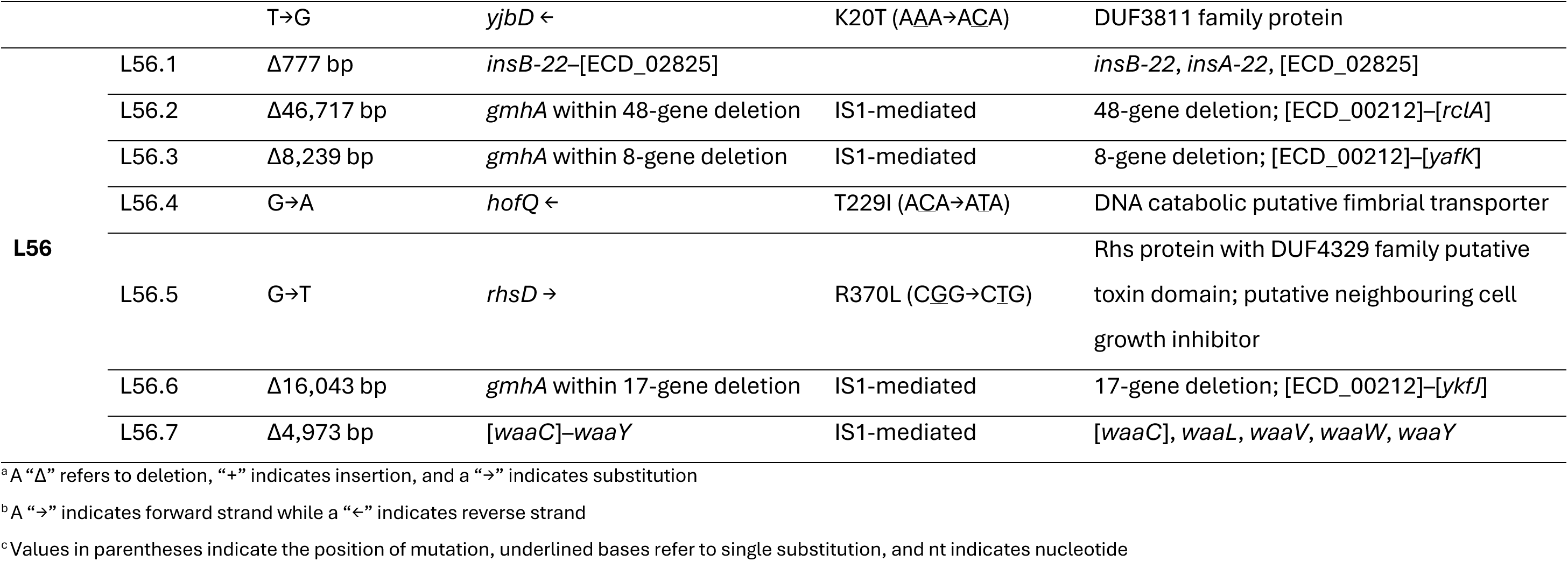
Phage-resistant mutations against evolutionary trained BLS5.

### Supplementary Figures

**Supplementary Figure 1:**
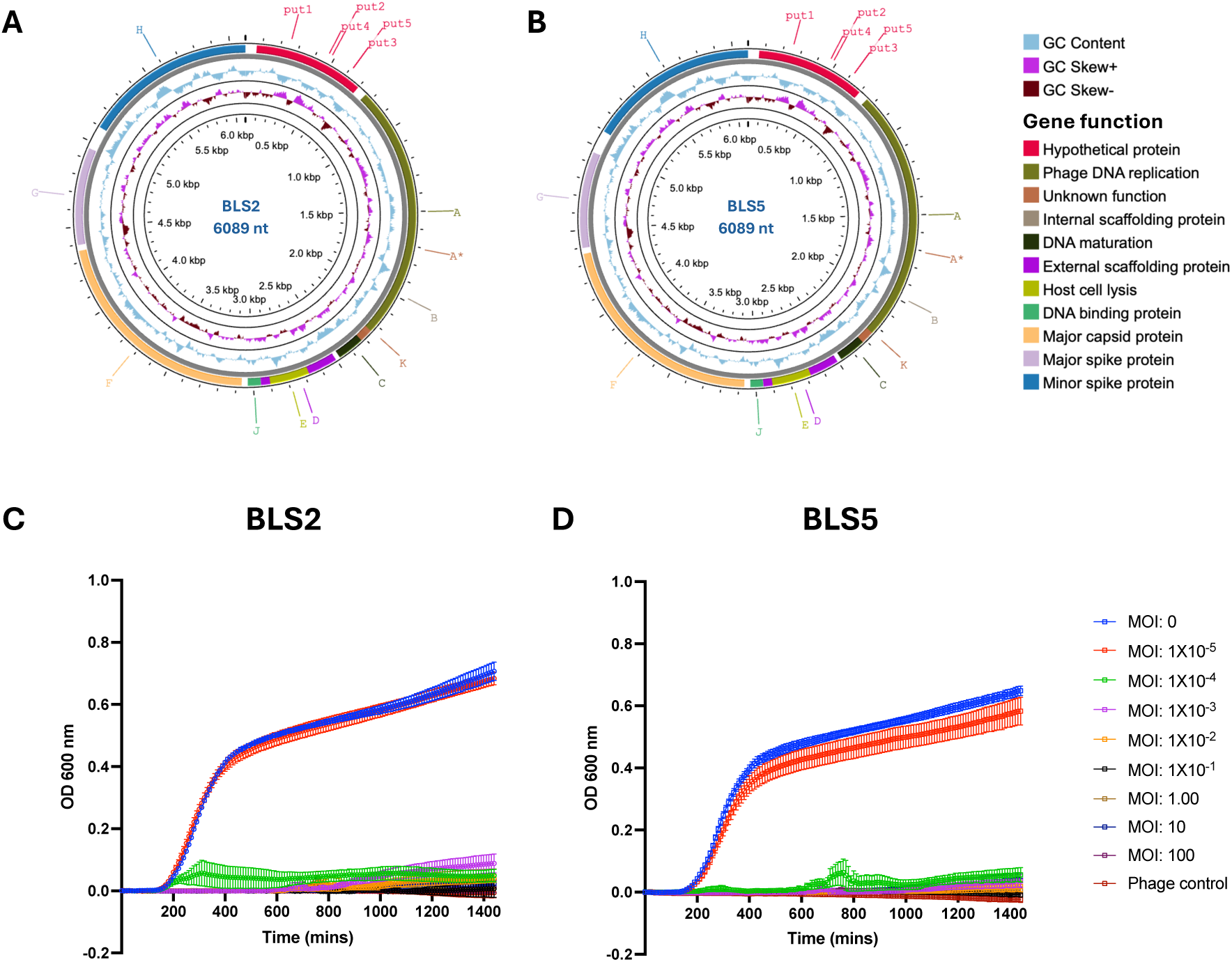
Characterisation of phages BLS2 and BLS5. Representation of draft genomes of **A)** BLS2 and **B)** BLS5. The outermost circle represents the phage genes. The middle circle represents GC content, and the innermost circle represents GC skew. Growth kinetics of *E. coli* BL21 in the presence of **C)** BLS2 and **D)** BLS5 at varying MOIs (Supplementary Data 1C-D).

**Supplementary Figure 2:**
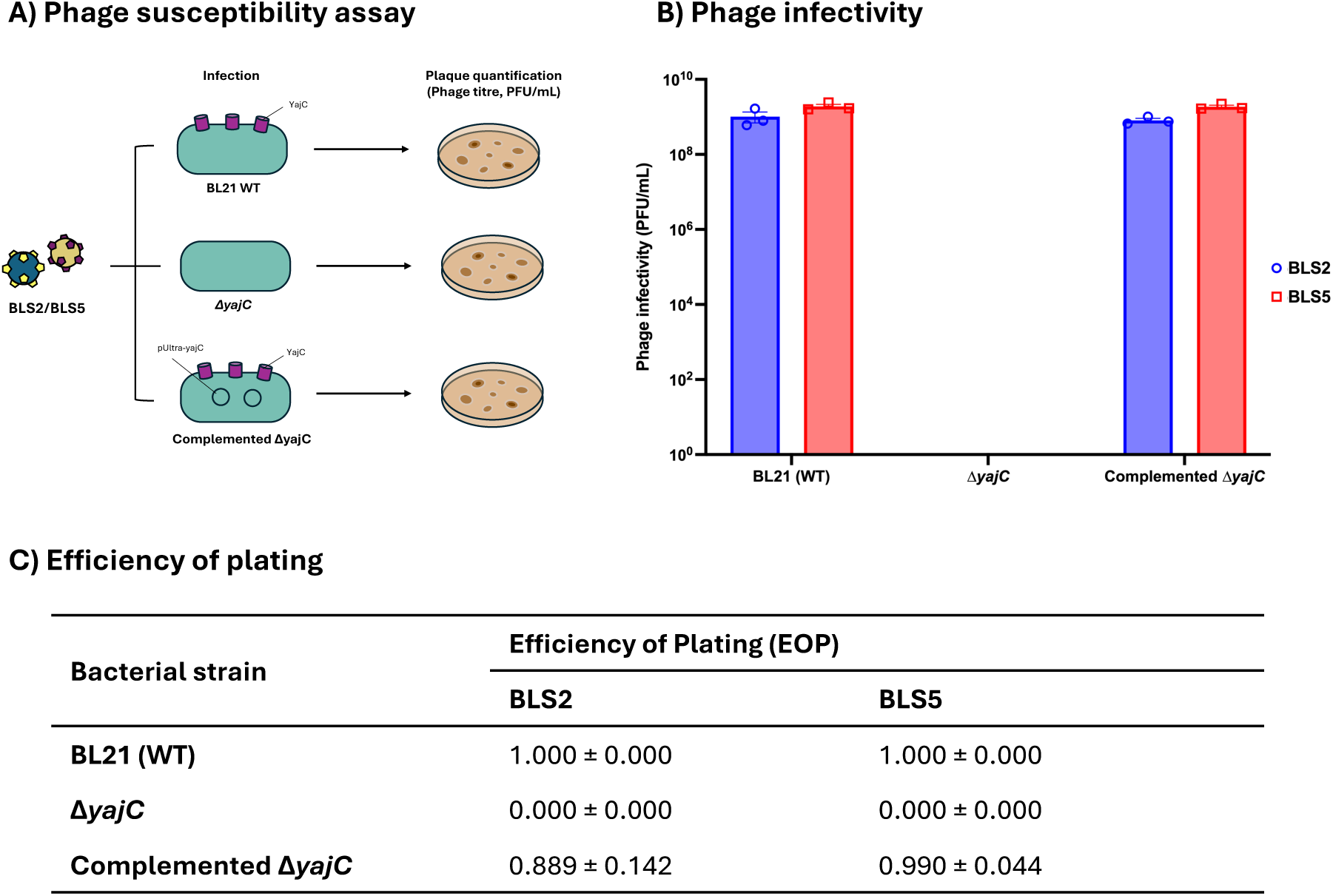
Susceptibility of *E. coli* BL21, Δ*yajC* and complemented Δ*yajC* to phages BLS2 and BLS5. **A)** Overview of phage susceptibility assay. Bacteria were infected with phages, and the number of plaques was quantified using the double agar overlay assay. **B)** The infectivity of phages BLS2 and LS5 against *E. coli* BL21, Δ*yajC,* and complemented Δ*yajC*. **C)** The efficiency of plating for BLS2 and BLS5. Phages that produce an equal number of plaques as the host *E. coli* BL21 have an EOP of 1.000. All data were presented as mean ± standard error.

**Supplementary Figure 3:**
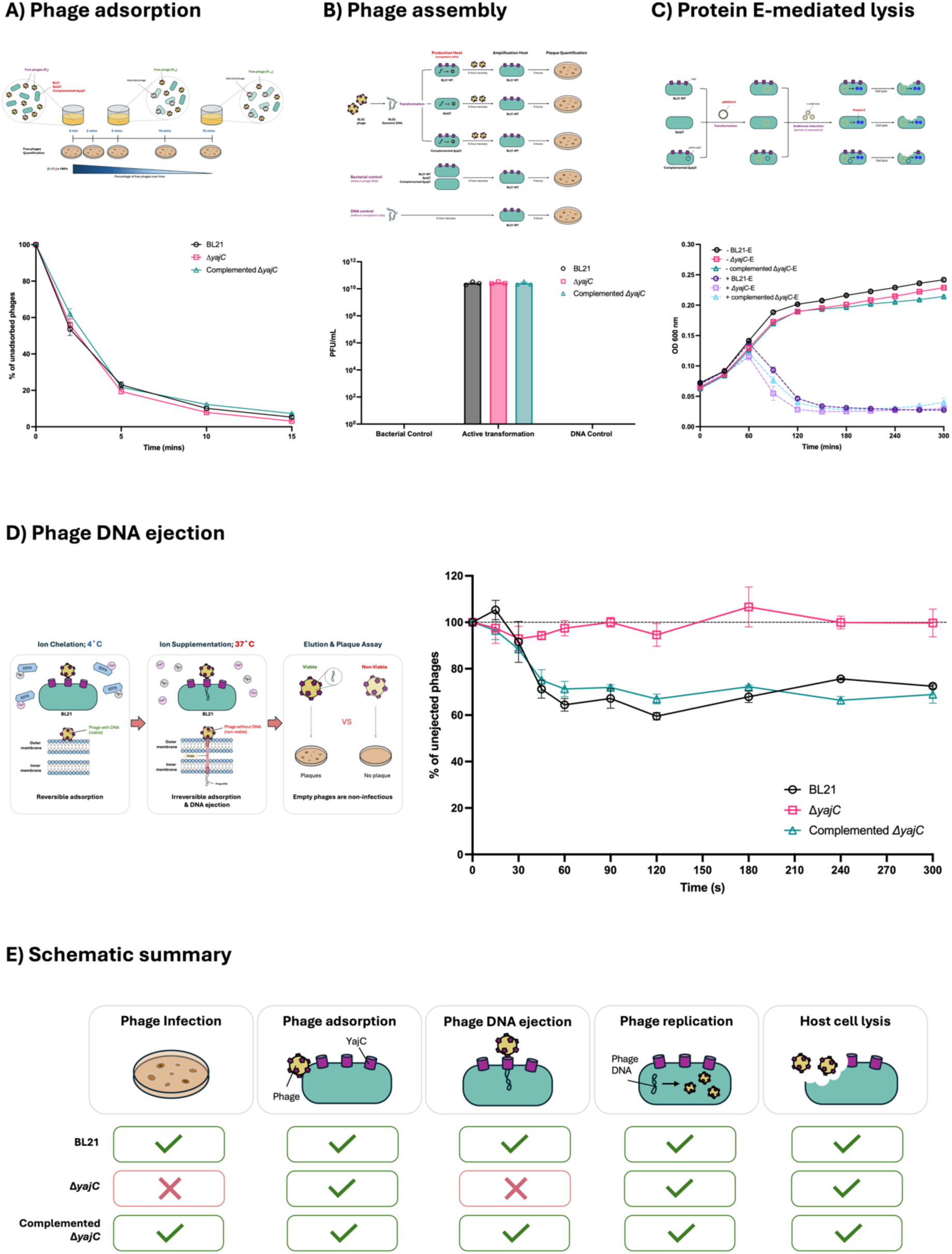
YajC is required for phage genome translocation. **A)** Adsorption of phage BLS2 on *E. coli* BL21, Δ*yajC* and complemented Δ*yajC*. Phages at a known titer (P_0_) were added to exponentially growing bacteria. Aliquots were taken at specific timepoints to determine the number of free phages (P_T_). **B)** Phage replication and virion assembly assay. Phage DNA was transformed into bacteria and recovered for two hours at 37°C to allow for phage replication and production. The lysate was collected and incubated with *E. coli* BL21 to amplify the phages, and the number of plaques was quantified using the double agar overlay assay. Negative controls without phage DNA or competent cells were included. Plaque formation (PFU/mL) after transformation of BLS2 genomic DNA into chemically competent *E. coli* BL21, Δ*yajC* and complemented Δ*yajC* via heat-shock. **C)** Gene *E*-mediated cell lysis assay. Gene *E* was induced using L-arabinose, and the bacterial growth was monitored over time. Growth kinetics of *E. coli* BL21, Δ*yajC*, and complemented Δ*yajC* upon gene *E* induction. “+” indicates L-arabinose induction; “-” indicates no induction. **D)** Phage genome ejection assay. Phages were first allowed to adsorb onto the host reversibly at 4°C and under ion-chelated conditions. Divalent cations were then supplemented, and the temperature was raised to 37°C to promote irreversible binding and genome ejection. Given that ejected phages are non-viable, they are unable to infect, hence no plaque formation. Phage genome ejection kinetics over the first five minutes, in *E. coli* BL21, Δ*yajC* and complemented Δ*yajC*, expressed as the percentage of phages that have not ejected their genomes. All data were presented as mean ± standard error. **E)** Schematic summary of the role of YajC during *Microviridae* infection. Tick marks indicate unaffected processes, while crosses indicate impaired processes.

**Supplementary Figure 4:**
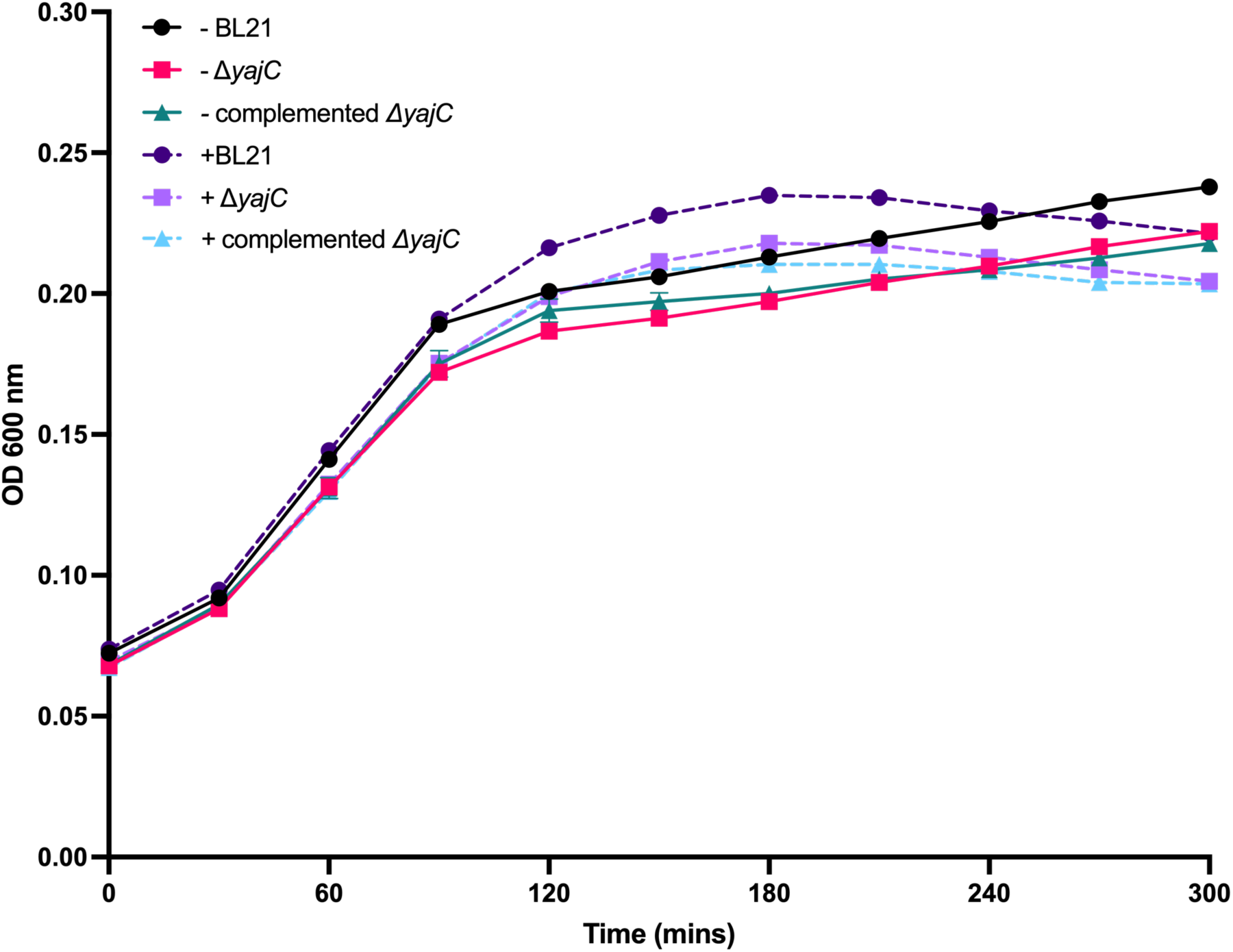
Growth kinetics of *E. coli* BL21, Δ*yajC*, and complemented Δ*yajC* upon induction of the pBAD24 vector. All data were presented as mean ± standard error. “+” indicates L-arabinose induction; “-” indicates no induction.

**Supplementary Figure 5:**
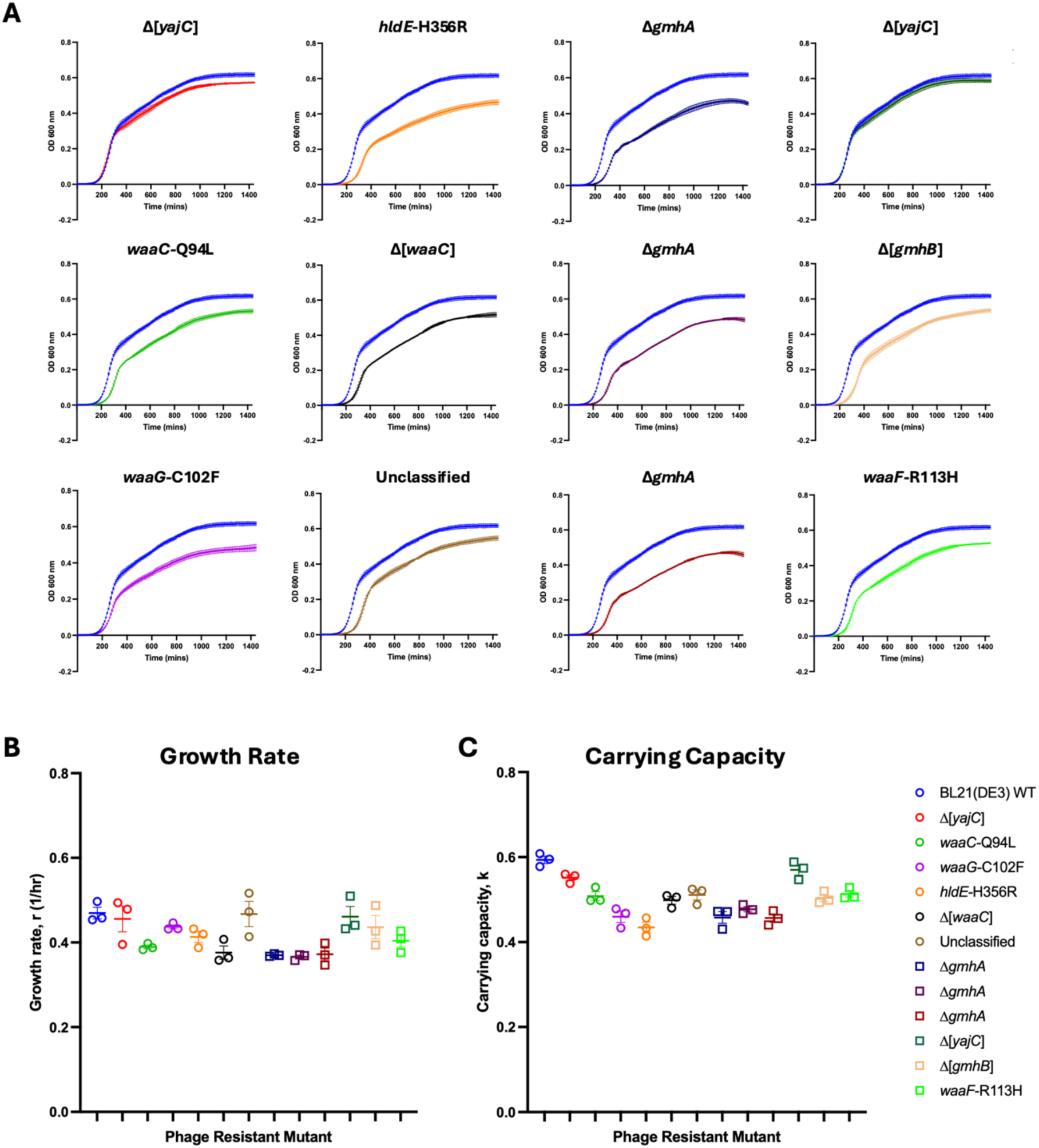
Fitness of phage-resistant bacteria. **A)** Growth curves, **B)** growth rates and **C)** carrying capacity of *E. coli* BL21 (blue) and its phage-resistant bacteria. Data is represented in mean ± standard error.

**Supplementary Figure 6:**
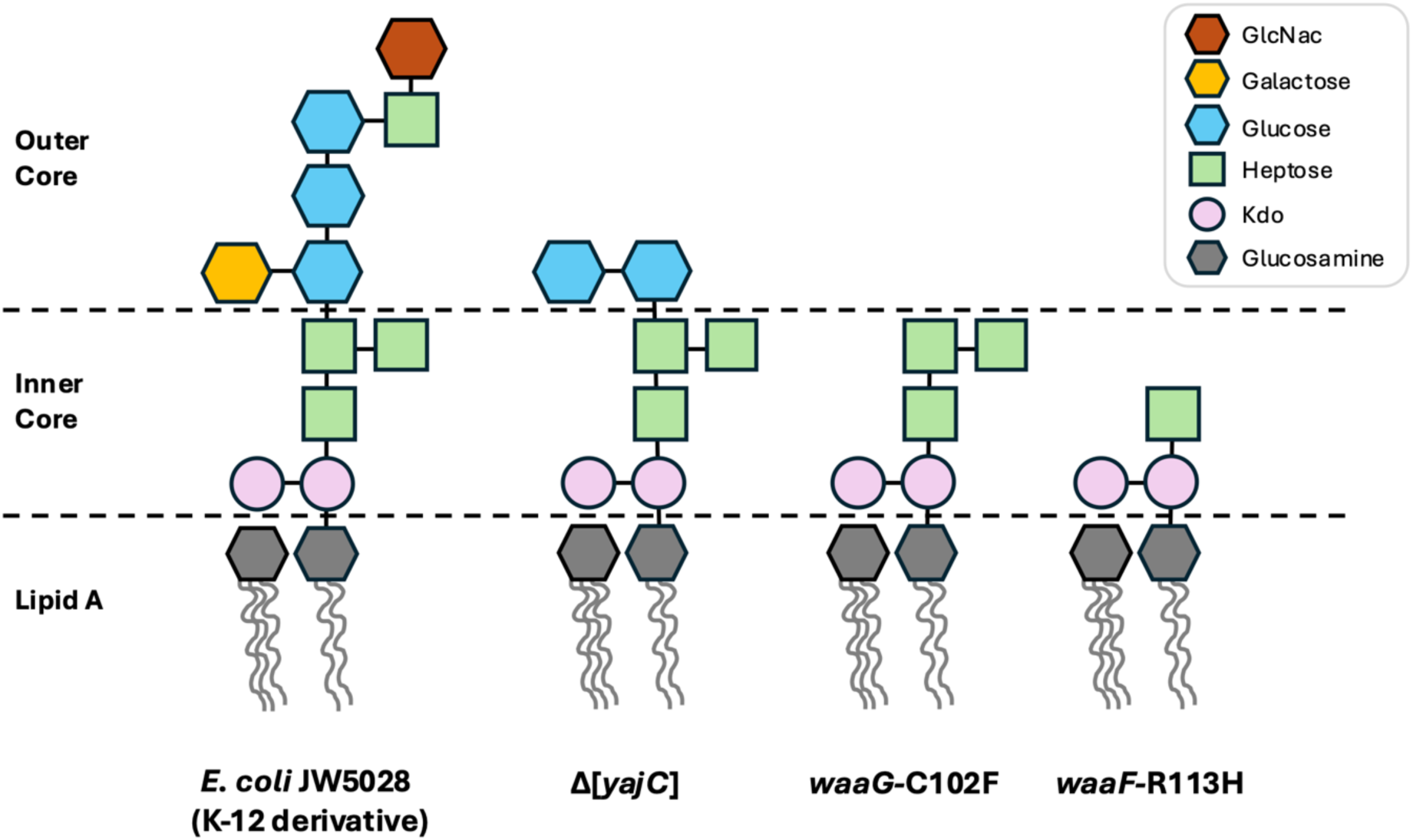
Schematic prediction of LPS profiles for the E. coli strains used in directed phage training (*2*). Prediction for Δ[*yajC*], *waaG*-C102F and *waaF*-R113H was made based on the functions of the mutated genes. Abbreviations in the diagram: GlcNac refers to N-acetylglucosamine, and Kdo refers to 3-deoxy-D-manno-oct-2-ulosonic acid.

**Supplementary Figure 7:**
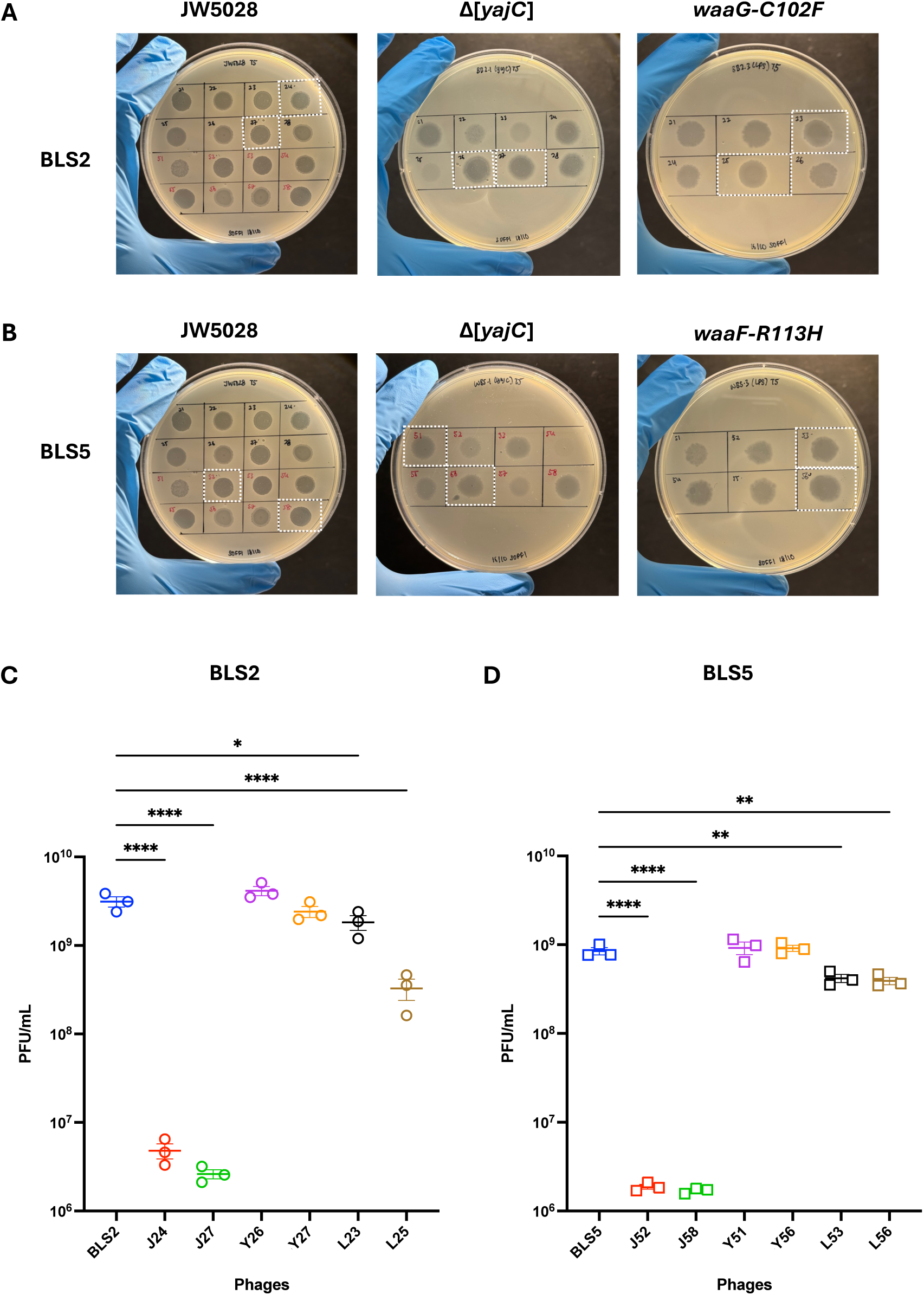
Infectivity of evolutionary trained phages. Representative images of spot assay from the last evolution cycle for **A)** BLS2 and **B)** BLS5. White boxes indicated evolutionarily trained phage populations selected for downstream analysis. **C-D)** Infectivity (PFU/mL) of evolutionary trained phages against *E. coli* BL21. Statistical significance was determined using One-way ANOVA, and a post-hoc Dunnett’s test was done to compare each trained phage to the ancestral phage. Asterisks denote the following significance levels: p < 0.05 (*), p < 0.01 (**), p < 0.001 (***) and p < 0.0001 (****).

## Supplementary Note

### Supplementary Note 1: YajC is required for phage infection

Microviridae infection comprises two kinetic stages. To initiate an infection, microviruses will first adsorb to the bacterial LPS. Given that they are tail-less phages, upon adsorption, microviruses may then “roll” along the cell surface to locate the second receptor, resulting in irreversible attachment and DNA ejection (*3*). Since the deletion of *yajC* confers phage resistance, we further investigated the role of YajC in *Microviridae* infection. This inner membrane protein was found to be associated with the injection of *Salmonella* phage P22 DNA into the bacterial host (*4*). Therefore, we hypothesised that YajC plays a similar role, serving as a second critical receptor, facilitating the translocation of *Microviridae* DNA during infection. To test this hypothesis, we examined whether YajC influences the remaining key infection stages, including phage adsorption, intracellular replication and assembly, and subsequent host cell lysis. Given that phages BLS2 and BLS5 belong to the same genus and share high genomic similarity (99.95%). As proof of concept, additional experiments were performed using phage BLS2 as a representative.

In the adsorption assay, over 80% of the phages were adsorbed to either *E. coli* BL21, Δ*yajC* or plasmid-complemented strain within the first ten minutes (Fig. S3A). Thus, YajC did not affect phage adsorption.

Next, we sought to determine if the deletion of *yajC* would affect phage DNA replication and virion assembly. To do so, we transformed BLS2 phage DNA directly into *E. coli* BL21, Δ*yajC* and complemented strains, bypassing the adsorption and DNA ejection stages. This enables us to evaluate whether the bacteria lacking YajC can replicate phage DNA and produce viable phage particles. We observed plaque formation in *E. coli* BL21 following phage DNA transformation, achieving a phage titer of 10^10^ PFU/mL. Similar titers were also obtained from Δ*yajC*, and upon complementation (Fig. S3B). These findings indicate that YajC is not part of the host replicative machinery because its deletion does not affect phage DNA replication and particle assembly.

Once phages are assembled, microviruses will lyse the host to release viral progeny. This process is mediated by the phage-encoded lysis protein E (*3*). Protein E binds and inhibits MraY, an essential integral membrane enzyme for peptidoglycan synthesis. Inhibition of MraY disrupts cell wall synthesis, ultimately compromising membrane integrity, leading to cell lysis (*5*). Given that the expression of protein E alone is sufficient to trigger bacterial lysis (*6, 7*), we introduced the plasmid carrying a wild-type copy of gene *E* into *E. coli* BL21, Δ*yajC*, and the complemented strain (Fig. S3C). Upon gene *E* induction, the optical density of *E. coli* BL21 decreased, indicating cell lysis. In contrast, no drastic change occurred when the empty pBAD24 vector was induced (Fig. S4), inferring that the reduction in cell density was in fact due to protein E activity. Similar lysis profiles were observed in Δ*yajC* and complemented Δ*yajC*. (Fig. S3C), indicating that YajC does not affect the efficiency of protein E-mediated cell lysis.

